# Inverse blebs operate as hydraulic pumps during mouse blastocyst formation

**DOI:** 10.1101/2023.05.03.539105

**Authors:** Markus F. Schliffka, Julien G. Dumortier, Diane Pelzer, Arghyadip Mukherjee, Jean-Léon Maître

**Affiliations:** Institut Curie, CNRS UMR3215, INSERM U934, PSL Research University, Sorbonne Université, Paris, France; Carl Zeiss SAS, Marly-le-Roy, France; Laboratoire de physique de l’École Normale Supérieure, CNRS UMR 8023, PSL Research University, Sorbonne Université, Université Paris Cité, Paris 75005, France

## Abstract

During preimplantation development, mouse embryos form a fluid-filled lumen, which sets their first axis of symmetry^1,2^. Pressurized fluid breaks open cell-cell contacts and accumulates into pockets, which gradually coarsen into a single lumen^3–5^. During coarsening, the adhesive and contractile properties of cells are thought to guide intercellular fluid (IF) but what cell behavior may control fluid movements is unknown. Here, we report large fluid-filled spherical membrane intrusions called inverse blebs^6,7^ growing into cells at adhesive contacts. At the onset of lumen coarsening, we observed hundreds of inverse blebs throughout the embryo, each dynamically filling with IF and retracting within a minute. We find that inverse blebs grow due to pressure build-up resulting from luminal fluid accumulation and cell-cell adhesion, which locally confines fluid. Inverse blebs then retract due to actomyosin contraction, which effectively redistributes fluid within the intercellular space. Importantly, inverse blebs show topological specificity and only occur at contacts between two cells, not at contacts formed by multiple cells, which essentially serve as fluid sinks. Manipulating the topology of the embryo reveals that, in the absence of sinks, inverse blebs pump fluid into one another in a futile cycle. We propose that inverse blebs operate as hydraulic pumps to promote luminal coarsening, thereby constituting an instrument used by cells to control fluid movement.

---

All living cells are immersed in fluid^8^. Within animal tissues, the amount of fluid between cells varies greatly from blood, where plasma constitutes the majority of tissue volume, to dense tissues, where IF is hardly detected. During embryonic development or the formation of organoids, relocation of IF is suspected to influence tissue jamming^9–11^ or topological transitions when fluid lumens form^12^. In some of the most striking relocations of IF, pressurized fluid can even fracture tissues^3,13,14^. In large fluid compartments, such as lumens, the movement of IF can be controlled by specialized cell protrusions called cilia^15^, but in packed tissues we are only beginning to understand how cells control the fluid surrounding them^16^.

During the formation of the first mammalian lumen, pressurized fluid fractures cell-cell contacts into hundreds of microlumens, which eventually coarsen into a single lumen called the blastocoel^3^ (Fig. 1A). As observed in mouse embryos^3^ and in a reconstituted in vitro model^17^, fracturing follows the path of lowest cell-cell adhesion and, once trapped into microlumens, fluid follows gradients of pressure set by cell mechanics. Therefore, the physical properties of cells guide IF throughout the embryo and determine the position of the first mammalian lumen, which is key to set its dorsoventral axis^1,2^. How active cellular processes modulate pressure gradients and guide IF in a complex three-dimensional geometry remains unknown. Regulation of actomyosin activity is a likely mechanism to modulate hydrostatic pressure locally^4,18–20^ and thus IF movements. However, the levels of actomyosin contractility are weak at cell-cell contacts, which translates into low tensions^3,21^. Moreover, actomyosin processes typically act on faster timescales (tens of seconds to minutes^22,23^) compared to the lifetime of microlumen networks (several hours)^3^. How cell contractility could act at cell-cell contacts is thus unclear.

**Figure 1:**
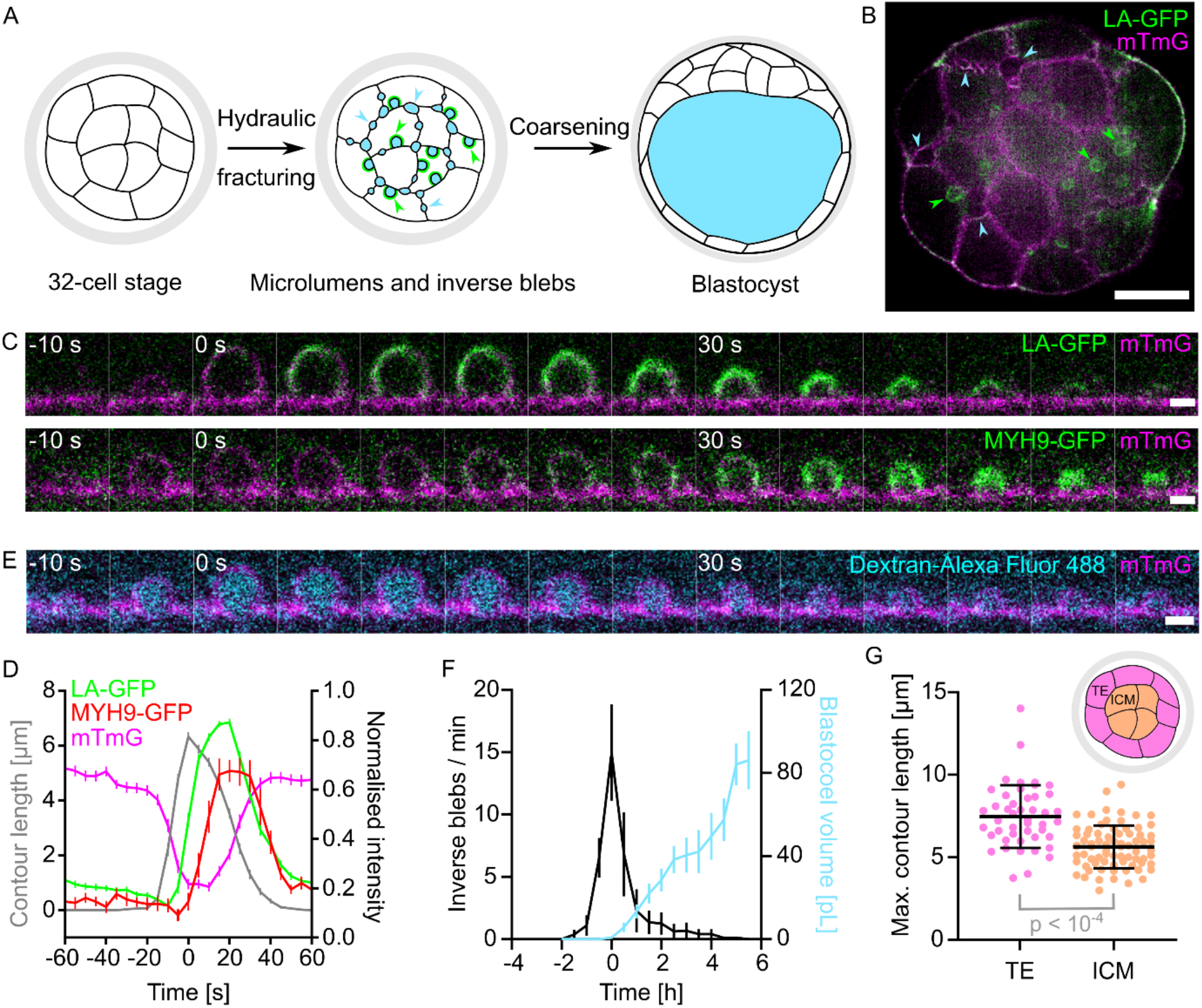
Inverse blebs at the onset of blastocoel formation. A) Schematic diagram of blastocyst lumen formation showing a 32-cell stage embryo before (left) and after (middle) hydraulic fracturing of cell-cell contacts and after the coarsening of fluid-filled (blue) pockets into a single lumen to form a blastocyst (right). After fracturing and before coarsening of the lumen, two types of fluid pockets can be observed: microlumens (blue arrowheads, symmetric, without actin in green) and inverse blebs (green arrowheads, asymmetric, coated with actin). B) Equatorial plane of a 32-cell stage mouse embryo with LifeAct-GFP (LA-GFP) in green and mTmG in magenta (Movie 1). Green arrowheads point at LA-GFP positive inverse blebs and blue arrowheads at LA-GFP negative microlumens. Scale bar, 20 μm. C) Still images following the lifetime of an inverse bleb shown with mTmG in magenta and LA-GFP (top, Movie 1) or MYH9-GFP (bottom, Movie 2) in green. Images taken every 5 s with time indicated relative to the maximal inverse bleb size. Scale bars, 2 μm. D) Mean contour length (grey, n = 121 blebs from 18 embryos) and normalized LA-GFP (green, n = 100 blebs from 13 embryos), MYH9-GFP (red, n = 21 blebs from 5 embryos) or mTmG (magenta, n = 142 blebs from 23 embryos) intensity of inverse blebs synchronized to their maximal extension. Error bars show SEM. E) Still images following the lifetime of an inverse bleb shown with Dextran-Alexa Fluor 488 (labelling intercellular fluid) in cyan and mTmG in magenta (Movie 3). Images taken every 5 s with time indicated relative to the maximal inverse bleb size. Scale bar, 2 μm. F) Number of inverse blebs per minute observed in embryos imaged every 10 s for 5 min repeated every 30 min in their full volume using light-sheet microscopy (Movie 4) during blastocoel formation. The volume of the segmented blastocoel is shown in blue. Mean ± SEM values of 8 embryos are calculated after synchronizing embryos to the time of maximal inverse bleb number. G) Maximal contour length of inverse blebs growing in trophectoderm (TE, pink, n = 46 blebs from 18 embryos) and inner cell mass (ICM, peach, n = 75 blebs from 18 embryos) cells. Mean and SD are shown in black bracket. p value results from Student’s t test. Top-left: schematic diagram showing TE and ICM cells location in a 32-cell stage embryo.

To examine whether fast actomyosin processes occur during lumen positioning, we imaged 32-cell stage mouse embryos at the onset of lumen formation with high spatiotemporal resolution. Imaging the plasma membrane, we observed short-lived hemispherical intrusions at cell-cell contacts growing several microns into cells in ∼15 s before retracting within ∼40 s (Fig. 1B-D, Movie 1-2). Compared to microlumens growing ∼1 μm for ∼3 h between cell-cell contacts^3^, such intrusions are larger and shorter lived. Actin and non-muscle myosin II appear at these intrusions only as they retract, with actin detected shortly before reaching maximal size and myosin arriving ∼5 s later (Fig. 1D and Supplementary Fig. 1A, Movie 1-2). Both actin and myosin signals increase, rapidly peaking ∼20 s after the maximal extension of the intrusion, before they return to the weak basal level typical of cell-cell contacts^21^ (Fig. 1A-C, Supplementary Fig. 1B, Movie 1-2). Such dynamics are characteristic of blebs, which are hemispherical protrusions growing without actin before being retracted by actomyosin within ∼1 min^20,24,25^. Therefore, we will refer to these intrusions as *inverse blebs*, since contrary to classical outward blebs they grow into the cytoplasm. Inverse blebs have been described previously at the free apical surface of zebrafish endothelial cells^6^ and of mouse hepatocytes^7^. Here, inverse blebs appear at basolateral cell-cell contacts. Importantly, inverse blebs forming at cell-cell contacts of mouse embryos do not result from a cell pushing an outward bleb into its neighbor. Rather, we observed decreased membrane signal intensity at the inverse bleb surface (Fig. 1D), indicating that plasma membranes from contacting cells locally detach and accumulate IF, as confirmed by labelling IF with a fluorescent dextran (Fig. 1E, Movie 3). Also, the intrusions are not secretory vesicles^26,27^ since membrane could clearly be seen expanding from the cell-cell contact while IF fills them up (Fig. 1C, E), Movie 1-3). We also considered the possibility that intrusions would be macropinocytotic cups, which also fill up with IF^28^. However, macropinocytotic cups grow due to actin polymerization, which is not the case with the intrusions observed here expanding without actin (Fig. 1C-E, Movie 1-2). Taken together, these observations reveal inverse blebs operating at very fast timescales, inflating and retracting within a minute at the adhesive contacts of mouse embryos (Fig. 1D).

To investigate the link between short-lived inverse blebs and long-term blastocoel formation, we performed full volume light-sheet microscopy to image inverse blebs throughout lumen formation (Movie 4) and quantified their spatiotemporal dynamics. Inverse blebs can be observed for about 3 h during lumen formation, with a maximum of ∼15 blebs per minute on average (Fig. 1F). The maximal blebbing activity precedes the rapid growth of the lumen. In fact, by the time the blastocoel reaches a size of ∼5 pL, equivalent to the volume of a single 32-cell stage blastomere^3,29^, over 80 % of blebs have already occurred (Fig. 1F, Supplementary Fig. 2A). Therefore, inverse blebs occur transiently during the nucleation phase of lumen formation. Using spinning disk confocal microscopy to resolve smaller microlumens, we systematically observed microlumens appearing ∼20 min before the first inverse bleb and persisting ∼90 min after the last inverse bleb (Supplementary Fig. 2B, Movie 5). Therefore, short-lived inverse blebs do not precede microlumens, but always coexist with them and share the same IF to inflate (Movie 3). Microlumens and inverse blebs form at the interface of both trophectoderm (TE) and inner cell mass (ICM) cells, the two cell lineages forming the blastocyst. Previously, we found that due to the different mechanics of TE and ICM cells, microlumens show distinct shapes at their respective interfaces^3^. Here, we measured larger inverse blebs in TE than in ICM cells (Fig. 1G, Supplementary Fig. 1C), which retract within the same time (Supplementary Fig. 1D). The different growth could be explained by the lower pressure in TE cells compared to ICM cells^3,30^ and indicates that an inverse bleb growing in TE cells displaces more fluid than when growing in ICM cells. We further observed that inverse blebs appear without significant bias at all types of cell-cell contacts: between two TE cells, two ICM cells or between a TE and an ICM cell (once corrected for contact type sampling, Supplementary Fig. 2C-F). Finally, counting inverse blebs throughout lumen formation and analyzing their position relative to the final location of the blastocoel revealed no clear relationship (Supplementary Fig. 2G). Taken together, we find that inverse blebs are temporally regulated and show lineage specific characteristics.

Unlike outward blebs and other inverse blebs observed so far, inverse blebs of preimplantation embryos grow at adhesive cell-cell contacts. On the one hand, adhesion could prevent the nucleation and growth of inverse blebs by keeping the intercellular space shut. On the other hand, adhesion molecules form clusters at the edges of microlumens^3^ and at the neck of inverse blebs (Supplementary Fig. 3A-D, Movie 6), and thus could trap the fluid needed for the growth of inverse blebs. To explore the role of adhesion, we imaged embryos in which *Cdh1* is maternally knocked out (m*Cdh1*^+/-^)^31^. m*Cdh1*^+/-^ embryos form a blastocyst while displaying microlumens for a significantly shorter time than wildtype (WT) embryos (Supplementary Fig. 4A-B, Movie 7). In m*Cdh1*^+/-^ embryos, we observed that microlumens tend to fuse into a single lumen instead of progressively coarsening by exchanging fluid at a distance as observed in WT embryos (Movie 7). This is consistent with cell-cell contacts being less adhesive and providing less mechanical resistance to the accumulation of pressurized fluid. Hence, reduced adhesion increases hydraulic compliance within the embryo, as evidenced in reconstituted in vitro settings^17^. In m*Cdh1*^+/-^ embryos, we rarely observed inverse blebs and, if so, in reduced numbers compared to WT embryos (Fig. 2A-B, Supplementary Fig. 4C, Movie 8), which, as suggested above, indicates a critical role of adhesion in entrapment of fluid at cell-cell contacts needed for inverse bleb nucleation and/or growth. Since adhesion could promote the local accumulation of fluid in pockets of low adhesion, we tested the local effects of adhesion by generating chimeric embryos with half of the embryo expressing lower levels of CDH1. This was achieved by knocking out *Cdh1* zygotically (z*Cdh1*) via CRISPR/Cas9 microinjection at the 2-cell stage (Supplementary Fig. 4D-E). As reported previously^3^, chimeric embryos fractured cell-cell contacts and formed a lumen preferentially located on the hydraulically compliant z*Cdh1* side of chimeric embryos (Supplementary Fig. 4E). In z*Cdh1* chimeric embryos, we did not observe inverse blebbing on either the mutant or WT side of the embryo (Supplementary Fig. 4F), revealing that a regional reduction of adhesion can alter the global hydraulics of the embryo. Therefore, adhesion provides an essential local confinement of intercellular fluid, which promotes inverse blebbing and can alter the mode of lumen coarsening globally.

**Figure 2:**
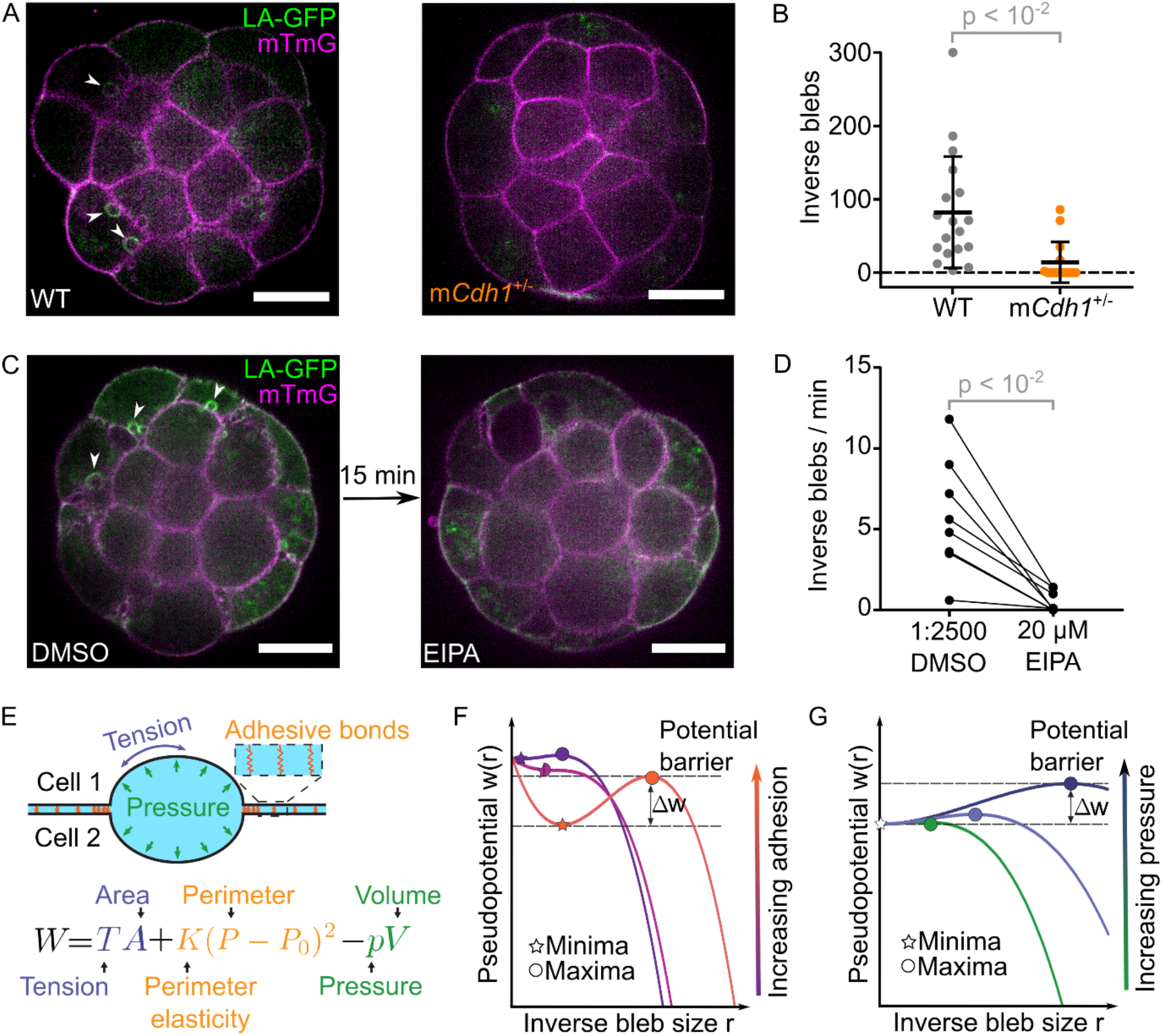
Inverse blebs formation requires cell-cell adhesion and fluid accumulation. A) Equatorial planes of WT (left) and m*Cdh1*^+/-^ (right) embryos with LifeAct-GFP (LA-GFP) in green and mTmG in magenta (Movie 7). Arrowheads point at inverse blebs. Scale bars, 20 μm. B) Number of inverse blebs observed at the equatorial plane of WT (grey, n = 18) and m*Cdh1*^+/-^ (orange, n = 15) embryos imaged every 1 min using spinning disk microscopy. Mean and SD are shown in black bracket. p value results from Mann-Whitney U test. C) Equatorial planes of embryos with LifeAct-GFP (LA-GFP) in green and mTmG in magenta in 1:2500 DMSO control medium and after 15 min of treatment with 20 μM EIPA (Movie 9). Arrowheads point at inverse blebs. Scale bars, 20 μm. D) Number of inverse blebs observed per minute in the equatorial plane of 8 embryos in 1:2500 DMSO control medium and after 15 min of treatment with 20 μM EIPA. p value results from paired Wilcoxon test. E) Schematic diagram of a fluid pocket trapped between two cells. The pseudopotential W characterizing the propensity of the fluid pocket to form an inverse bleb depends on the Tension T at the surface of the pocket of area A, the perimeter elasticity K of the adhesion bonds (orange) around the pocket of perimeter P and resting perimeter P_0_ and, the hydrostatic pressure p of the pocket of volume V. F-G) Pseudopotential w as a function of the inverse bleb size r with varying adhesion strength k (F) or pressure p (G). Normalized units, see Supplementary Note. For each example, the minima (star) and maxima (circle) are indicated. Only conditions with sufficient adhesion or low pressure allow for a finite potential barrier Δw needed to maintain the coexistence of microlumens and inverse blebs.

Outward blebs inflate because the cytoplasmic pressure is higher than the medium pressure, which results from the actomyosin cortex compressing the cytoplasm^19^. In the case of inverse blebs observed in zebrafish endothelial cells, inverse blebs inflate due to blood pressure^6^. In preimplantation embryos, we suspected that accumulation of luminal fluid could be responsible for the pressure increase necessary for inverse blebs. Luminal fluid accumulates by osmosis, to prevent this we blocked sodium transport by inhibiting sodium/proton exchangers with ethylisopropyl amiloride (EIPA) or the sodium/potassium pump with ouabain^32,33^. Either treatment prevented lumen formation in most embryos and inverse blebs could rarely be observed, and if so only when lumen formation would occur (Supplementary Fig. 5). To dynamically tune the accumulation of luminal fluid, we acutely treated blebbing embryos with EIPA, which caused inverse blebs to almost completely vanish within 15 min of treatment (Fig. 2C-D, Movie 9). Importantly, acutely treated embryos conserved their IF, microlumen network and small lumens that had formed before the treatment (Fig. 2C, Movie 9), indicating that there would be in principle enough fluid to inflate inverse blebs. Therefore, luminal fluid accumulation and the resulting build-up of hydrostatic pressure are required to inflate inverse blebs.

These initial observations hint at the following scenario: fluid accumulation, confined between adhesion pockets (Supplementary Fig. 3A-B), raises IF pressure until a first threshold is reached for microlumens to form (Supplementary Fig. 2B); further raising IF pressure above a second threshold nucleates inverse blebs, which stop when fluid ceases to accumulate (Fig. 2C-D) and does not need to occur in the absence of firm adhesion throughout the embryo (Fig. 2A-B, Supplementary Fig. 4C-F); finally, once a sufficiently large lumen of a few pL has formed (Fig. 1F, Supplementary Fig. 2A), adhesion confinement is locally reduced and IF pressure drops, leading first to the disappearance of inverse blebs and then of microlumens (Supplementary Fig. 2B). To better conceptualize the roles of cell-cell adhesion and fluid accumulation in the nucleation of inverse blebs, we formulate a minimal physical model of fluid pockets confined between two cells, which could form both stable microlumens or dynamic inverse blebs (Supplementary Note). Taking into account the pressure build-up within the IF, the elasticity provided by clustered adhesion sites at the periphery and tensions at the surface of fluid pockets (Fig. 2E, Supplementary Note), the mechanics of the system can be captured by a pseudopotential W (Fig. 2E-G). The minimum of the potential indicates mechanically stable fluid pockets with finite-sizes. This is also depicted in a recent theoretical framework associated to in vitro reconstruction of microlumen formation^17^. Here, we also find that beyond a critical size, symmetric lens-shaped microlumens go through a hydraulic instability^34^ leading to asymmetric hemispherical shapes, characteristic of inverse blebs (Supplementary Note). With sufficient adhesion, the model captures the coexistence of finite-sized and long-lived microlumens with dynamic inverse blebs (where pseudopotential is minimum at smaller sizes of fluid pockets in Fig. 2F). Here, the nucleation of inverse blebs is a potential barrier hopping phenomenon and lowering the potential barrier increases the susceptibility to nucleate inverse blebs. When adhesion is reduced, depending on the initial size, microlumens either shrink due to surface tension or grow to maximal size resulting in the absence of stable microlumens or inverse blebs (Supplementary Note), as observed when *Cdh1* is knocked out (Fig. 2A-B, Supplementary Fig. 4A-B). Similarly, the model suggests that more fluid accumulation and hence higher pressure build-up lowers the barrier (Fig. 2G). This is consistent with our experimental results where reducing fluid accumulation stops inverse bleb nucleation (Fig. 2C-D, Supplementary Fig. 5). Together, our theoretical framework qualitatively captures the effects of adhesion and pressure on the nucleation of inverse blebs that we observed experimentally.

Once inflated by pressurized luminal fluid, inverse blebs eventually stop growing and retract. To retract, outward blebs rely on the recruitment of actomyosin to contract the swollen protrusion^24^. To elucidate the role of actomyosin contractility on inverse blebs retraction, we used para-Nitroblebbistatin to acutely block all non-muscle myosin II heavy chains (NMHC) of blebbing mouse embryos. Inhibition of NMHCs increased the time taken to retract inverse blebs (Supplementary Fig. 6A-C, Movie 10), indicating that bleb retraction relies on actomyosin contractility. Among NMHC paralogs, outward bleb retraction is mediated by MYH9^35^, which we also detect at inverse blebs during their retraction (Fig. 1C-D, Supplementary Fig. 6J, Movie 2). We generated heterozygous mutants lacking the maternal allele of *Myh9* (m*Myh9*^+/-^), which are able to form a blastocyst^36^, and measured inverse bleb dynamics. Inverse blebs of m*Myh9*^+/-^ embryos grow as large as in WT embryos but take longer to retract (Fig. 3A-C, Supplementary Fig. 6G, Movie 11). In contrast, embryos lacking the maternal allele of *Myh10*, the other NMHC paralog expressed in the mouse preimplantation embryo^36^, did not show altered inverse bleb dynamics (Fig. 3A-C, Movie 11). Therefore, like outward blebs^35^, inverse blebs retract thanks to the recruitment of MYH9. Interestingly, inhibition of contractility did not affect actin recruitment dynamics or the growth and retraction profiles of inverse blebs (Fig. 3C, Supplementary Fig. 6C-I), unlike outward blebs, whose growth is impacted by reduced cytoplasmic pressure after contractility loss^19,35^. This further confirms our experimental and theoretical conclusions that inverse blebs inflate as a result of pressure from adhesive confinement and IF accumulation rather than from cell contractility (Fig. 2, Supplementary Note). In our theoretical framework, a time modulated increase of active tension, corresponding to the recruitment of actomyosin after inverse bleb extension, can change the underlying pseudopotential (Fig. 3D) and contract a growing inverse bleb. The resulting life cycle of inverse blebs can thus be depicted as an orbit in phase space of inverse bleb size and pseudopotential W, showing qualitative agreement between the model (Fig. 3D) and experimental data (Fig. 3C, Supplementary Fig. 6C).

**Figure 3:**
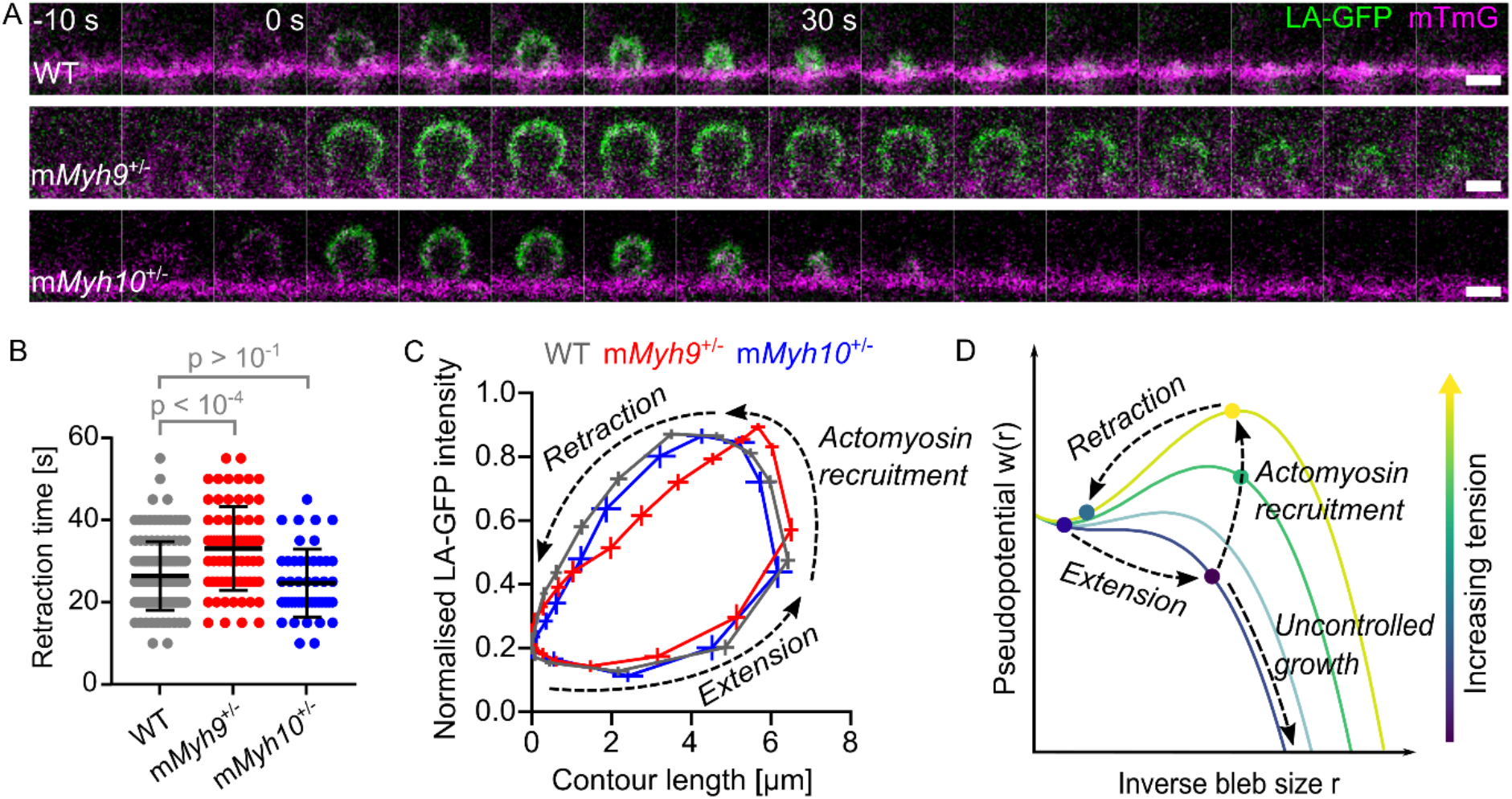
Inverse blebs retract with actomyosin contractility. A) Still images following the lifetime of an inverse blebs shown with LA-GFP in green and mTmG in magenta in WT (top), m*Myh9*^+/-^ (middle) and m*Myh10*^+/-^ (bottom) embryos (Movie 11). Images taken every 5 s with time indicated relative to the maximal inverse bleb size. Scale bars, 2 μm. B) Duration of retraction of inverse blebs for WT (grey, n = 121 from 18 embryos), m*Myh9*^+/-^ (red, n = 67 from 10 embryos) and m*Myh10*^+/-^ (blue, n = 41 from 6 embryos) embryos. Mean and SD are shown in black bracket. p values result from ANOVA and Dunnett’s tests. C) Mean normalized LA-GFP intensity as a function of mean contour length of inverse blebs synchronized to their maximal extension for WT (grey, n = 100 from 13 embryos), m*Myh9*^+/-^ (red, n = 67 from 10 embryos) and m*Myh10*^+/-^ (blue, n = 41 from 6 embryos) embryos. Error bars show SEM. Arrows indicate the extension and retraction dynamics of inverse blebs. D) Pseudopotential w as a function of the inverse bleb size r with varying tension. Normalized units, see Supplementary Note. Before actomyosin recruitment, fluid pockets undergo uncontrolled growth and extend into an inverse bleb. After actomyosin recruitment, the tension increases the pseudopotential w, stops the extension of the fluid pocket and eventually retracts it.

Due to their life cycle of fast fluid accumulation/charge and shrinkage/discharge dynamics, inverse blebs act as *bona fide* hydraulic pumps able to quickly redistribute fluid within the three-dimensional intercellular space. Dispersed inverse bleb fluid may fuel the formation of new blebs, which makes the effectiveness of such pumps to redistribute intercellular fluid questionable. However, the transport of this dispersed bleb fluid depends on the geometry of the intercellular space of the embryo. The intercellular space is made of three distinct topological entities: contacts between two cells or faces; tricellular contacts or edges; and multicellular junctions or vertices (Fig. 4A). Apart from their topology, these different types of contact sites likely differ vastly in their mechanics^3^. We had previously described two types of microlumens (Fig. 4A): bicellular microlumens, strictly confined between two cells (faces), and multicellular microlumens, which are surrounded by at least three cells (edges and vertices). Both bi- and multicellular microlumens grow and shrink during lumen formation but differ in size and dynamics^3^. We noted that inverse blebs form exclusively at the interface between two cells, both at bicellular microlumens and at seemingly closed cell-cell contacts, and do not appear at multicellular microlumens (Fig. 4A, Movie 1-3). This implies that multicellular microlumens situated at edges or vertices can behave as sinks, absorbing the fluid discharged by inverse blebs retraction. The presence of sinks makes inverse blebs effective hydraulic pumps, which could redistribute IF to promote the coarsening of the microlumen network.

**Figure 4:**
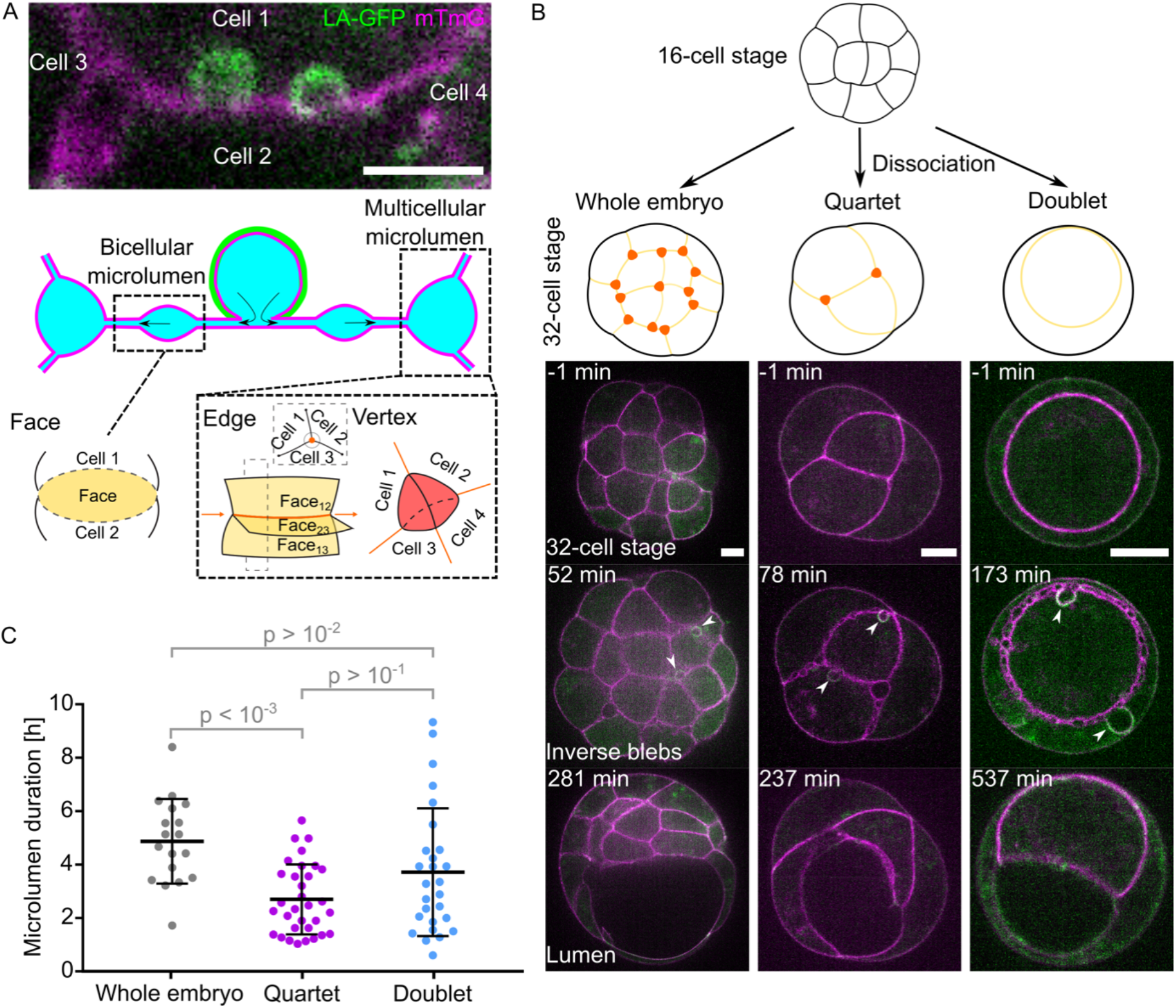
Inverse blebs pump fluid into topological sinks. A) Top: still image showing inverse blebs at a cell-cell contact bounded by two microlumens contained by multiple neighboring cells with LifeAct-GFP (LA-GFP) in green and mTmG in magenta. Middle: corresponding schematic diagram with cell membranes in magenta, actomyosin in green and IF in blue. Arrows indicate the fluid flows expected from the retraction of the inverse bleb. Bottom: Topological features associated with the microlumen network: contacts between two cells constitute a face and show bi-cellular microlumens and inverse blebs; at the contact of three or more faces respectively, edges and vertices show multicellular microlumens. B) Top: schematic diagram of a 16-cell stage embryo developing into a whole 32-cell stage embryo or into quartets and doublets after dissociation. Faces where bicellular microlumens form are shown in yellow. Edges and vertices where multicellular microlumens form are shown in red in whole embryos and quartets. Bottom: equatorial plane of a whole 32-cell stage embryo, quartet and doublet of 32-cell stage blastomeres with LA-GFP in green and mTmG in magenta (Movie 5, 12-13). Images are shown before microlumen appearance, during the presence of microlumens and inverse blebs and after their disappearance. Arrowheads point at inverse blebs. Scale bar, 10 μm. C) Microlumen duration during lumen formation in whole embryos (grey, n = 18 embryos), quartets (purple, n = 32 quartets) and doublets (blue, n = 27 doublets). Mean and SD are shown in black bracket. p values result from Kruskal-Wallis and Dunn’s multiple comparison tests.

To elucidate the interplay between inverse blebs and the topology of the intercellular space, we set out to modify the embryo geometry. In embryos with reduced cell number, there are fewer sinks that are also spatially located closer to each other compared to whole embryos. In this case, all discharged inverse bleb fluid should accumulate in these select nearby sinks, speeding up lumen formation. Contrarily, eliminating all such sinks would force all discharged inverse bleb fluid to circulate and nucleate new blebs, delaying the formation of the lumen. To test this idea, we dissociated embryos, creating doublets and quartets of 32-cell stage blastomeres to simplify the topology of the microlumen network (Fig. 4B, Supplementary Fig. 7A). With one eighth the cell number, quartets form microlumens within a smaller cell-cell contact network than whole embryos (Supplementary Fig. 7A-B). Topologically, quartets show a reduced number of channels and vertices (either 1 or 2 vertices), hence a reduced number of hydraulic sinks compared to whole embryos (Fig. 4B). We measured that microlumens persist twice as long in whole embryos as compared to quartets (Fig. 4C, Movie 5 and 12). Therefore, a simpler microlumen network with fewer sinks takes less time to coarsen. Following the same rationale, further reducing the cell number to a doublet, with only a single interface to coarsen microlumens at, should form a lumen faster than whole embryos and quartets. However, this is not the case (Fig. 4C, Movie 5 and 13). Unlike quartets, doublets do not have multicellular microlumens (Fig. 4B-C). Thus, all microlumens can form inverse blebs, pumping fluid back and forth between microlumens in a futile cycle (Movie 13). Altogether, we conclude that inverse blebs act as hydraulic pumps that flush luminal fluid trapped at cell-cell contacts into topological sinks to foster the coarsening of the microlumen network into a single lumen.

Outward blebs have been observed in a variety of situations and were proposed to help the clearance of apoptotic cells^37^, to act as signaling hubs^38^, migratory protrusions^39^, pressure valves^18^, or, in the case of inverse blebs, as facilitators of apical membrane expansion^6^. Here, we find that inverse blebs are able to repair the damages of pressurized fluid percolating the tissue by pumping luminal fluid away from cell-cell contacts. We had noted previously that bicellular microlumens disappear before the multicellular ones during blastocoel formation^3^. The hydraulic pumping of inverse blebs into topological sinks contributes to the coarsening of the bicellular microlumens into multicellular ones. Therefore, inverse blebs provide a cellular mechanism explaining how contractility, which is weak at cell-cell contacts, can position the first mammalian lumen and first axis of symmetry of the mammalian embryo. More generally, inverse blebs constitute a unique mechanism for cells to handle IF in dense tissues where beating cilia may underperform. As such, in other dense tissues where pressure is elevated like the liver, kidney, swollen lymph nodes or wounded skin^7,14,40,41^, inverse blebs could operate as hydraulic pumps redistributing the fluid permeating tissues.

## Acknowledgements

We thank the imaging platform of the Genetics and Developmental Biology Unit at the Institut Curie (PICT-IBiSA@BDD, member of the French National Research Infrastructure France-BioImaging ANR-10-INBS-04) for their outstanding support, in particular Olivier Leroy for his help writing Metamorph journals; the animal facility of the Institut Curie for their invaluable help. We thank Viventis Microscopy for custom Python scripts for the LS1 Live. We thank Maria Almonacic and Marie-Hélène Verlhac for generously sharing the Lap2b-GFP plasmid. We thank Yohanns Bellaïche, Carles Blanch-Mercader and members of the Maître lab for critical reading of the manuscript.

Research in the lab of J.-L.M. is supported by the Institut Curie, the Centre National de la Recherche Scientifique (CNRS), the Institut National de la Santé Et de la Recherche Médicale (INSERM), and is funded by grants from the Fondation Schlumberger pour l’Éducation et la Recherche via the Fondation pour la Recherche Médicale, the European Research Council Starting Grant ERC-2017-StG 757557, the Agence Nationale de la Recherche (ANR-21-CE13-0027-01), the European Molecular Biology Organization Young Investigator program (EMBO YIP), the INSERM transversal program Human Development Cell Atlas (HuDeCA), Paris Sciences Lettres (PSL) QLife (17-CONV-0005) grant and Labex DEEP (ANR-11-LABX-0044) which are part of the IDEX PSL (ANR-10-IDEX-0001-02). M.F.S. is funded by a Convention Industrielle de Formation pour la Recherche (No 1113 2019/0253) between the Agence Nationale de la Recherche and Carl Zeiss SAS; as well as La Ligue contre le cancer. M.F.S. thanks the support from La Fondation des Treilles. A.M. acknowledges funding from the QBio Junior Research Chair programme of the QBio initiative of ENS-PSL and the Parisante Campus.

## Author contributions

M.F.S., J.G.D. and D.P. performed experiments and prepared data for analyses. M.F.S. J.G.D. and J.-L.M. designed the project, analyzed the data. A.M. developed the theoretical analysis. M.F.S., A.M. and J.-L.M. wrote the manuscript. M.F.S. and J.-L.M. acquired funding.

---

**Supplementary Figure 1:**
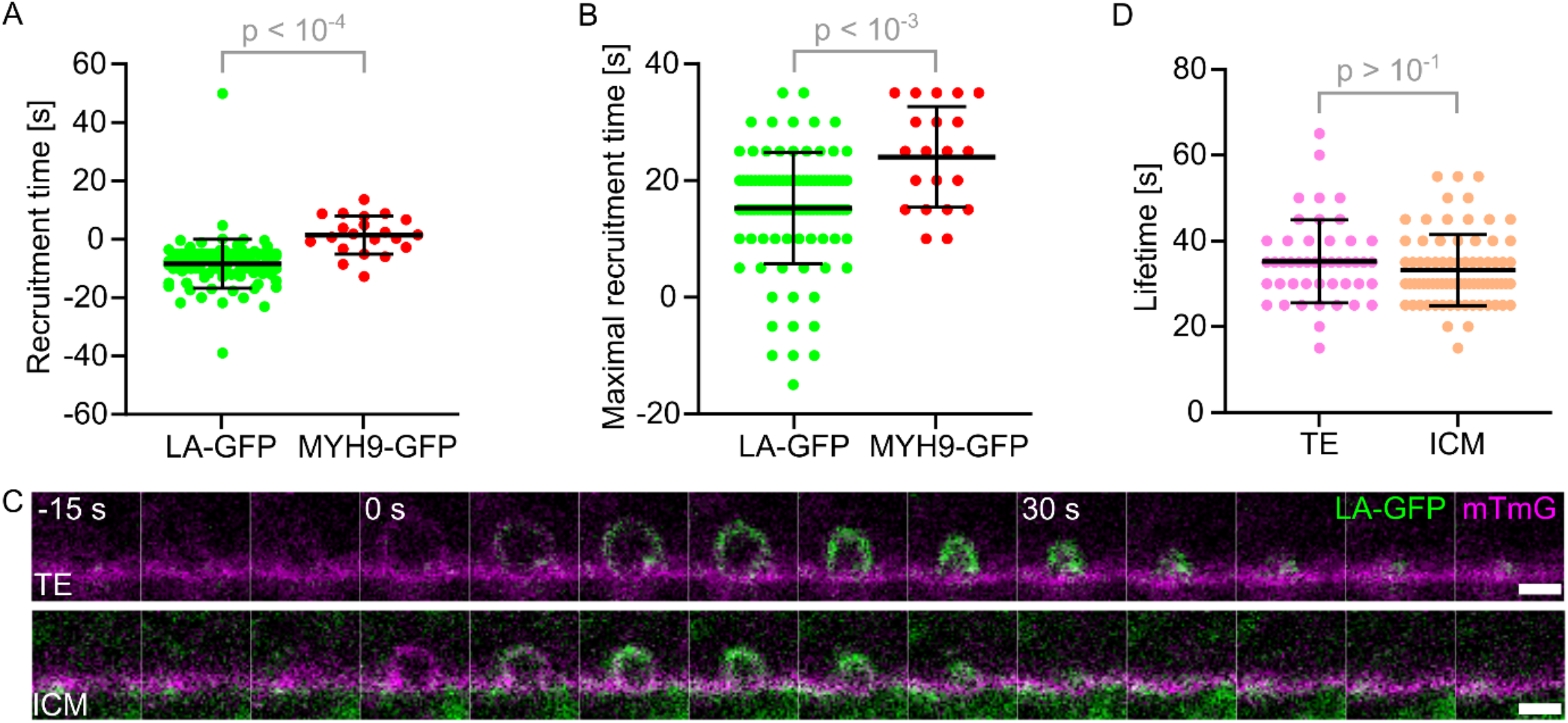
Dynamics of individual inverse blebs. A) Initial recruitment time of LA-GFP (green, n = 100 blebs from 13 embryos) and MYH9-GFP (red, n = 21 blebs from 5 embryos) to inverse blebs. Mean and SD are shown in black bracket. p value results from Student’s t test. B) Maximal recruitment time of LA-GFP (green, n = 100 blebs from 13 embryos) and MYH9-GFP (red, n = 21 blebs from 5 embryos) to inverse blebs. Mean and SD are shown in black bracket. p value results from Student’s t test. C) Still images following the lifetime of an inverse blebs shown with LA-GFP in green and mTmG in magenta for a TE (top lane) and an ICM (bottom lane) cell. Images taken every 5 s with time indicated relative to the maximal inverse bleb size. Scale bars, 2 μm. D) Inverse bleb lifetime in TE (pink, n = 46 blebs from 18 embryos) and ICM (peach, n = 75 blebs from 18 embryos) cells. Mean and SD are shown in black bracket. p value results from Student’s t test.

**Supplementary Figure 2:**
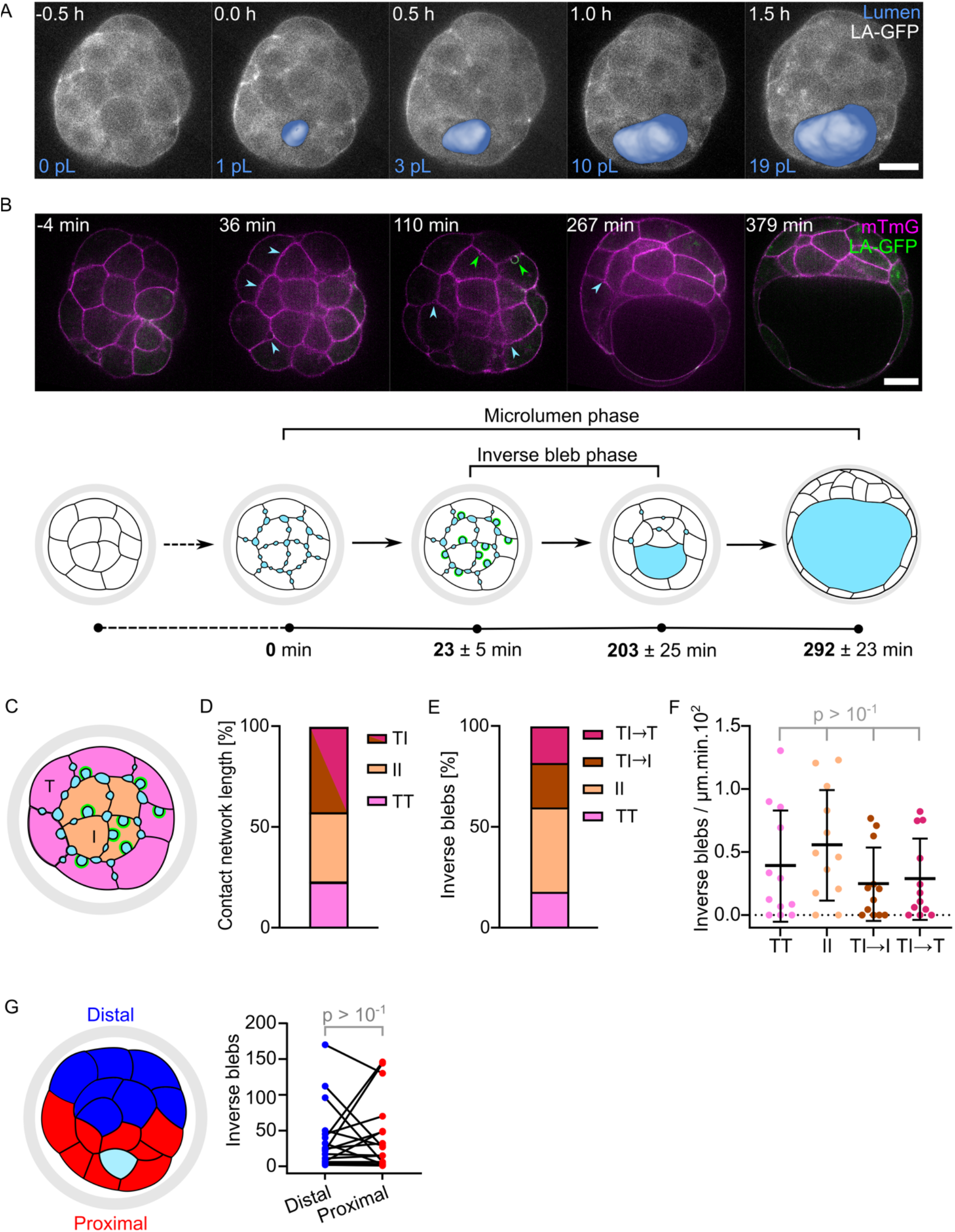
Spatiotemporal dynamics of inverse blebs throughout blastocoel formation. A) Equatorial plane of a 32-cell stage embryo with the segmented lumen (blue) overlaid on LA-GFP (grey, Movie 4). The volume of the segmented lumen is indicated on the bottom left. Scale bar, 20 μm. B) Top: equatorial plane of a 32-cell stage embryo with LA-GFP in green and mTmG in magenta (Movie 5) at distinct steps of lumen formation. Blue arrowheads point at LA-GFP negative microlumens, green arrowheads point at LA-GFP positive inverse blebs. Scale bar, 20 μm. Bottom: schematic diagrams of the distinct steps leading to the formation of the blastocoel (large blue structure) with mean ± SEM characteristic times measured in 18 embryos relative to the apparition of microlumens (small blue pockets) in a 32-cell stage embryo enclosed in the zona pellucida (grey). The microlumen phase corresponds to the time during which microlumens can be detected while the inverse bleb phase represents the duration of observation of inverse blebs (blue pockets coated with green actin). C) Schematic diagram of 32-cell stage embryo enclosed within the zona pellucida (grey) showing inverse blebs (asymmetric intrusions coated with green actomyosin and filled with blue luminal fluid) and microlumens (symmetric pockets of blue luminal fluid) between surface trophectoderm (pink, T) and inner cell mass cells (peach, I). D) Proportion of contact length relative to the cell-cell contact network within the imaging plane for TE-TE (TT, pink), ICM-ICM (II, peach) or TE-ICM (TI, brown-purple) from 12 embryos. E) Proportion of inverse blebs growing at TE-TE (pink), ICM-ICM (peach) and TE-ICM interfaces with the direction of growth into TE (brown) or ICM (purple) indicated for TE-ICM interfaces from 12 embryos. F) Number of inverse blebs observed per minute normalized by the length of cell-cell contacts at given tissue interfaces in the optical slice of embryos imaged using spinning disk microscopy (Movie 5). Inverse blebs are captured growing between trophectoderm cells (TT, n = 57), inner cell mass cells (II, n = 129), into inner cell mass cells (TI->I, n = 72) or into trophectoderm cells (TI->T, n = 63) at the interface between trophectoderm and inner cell mass cells of 12 embryos. Mean and SD are shown in black bracket. p values result from Kruskal-Wallis and Dunn’s multiple comparison tests. G) Left: schematic diagram of 32-cell stage embryo within the zona pellucida (grey) where cells were labeled as proximal (red) or distal (dark blue) relative to the position of the growing blastocoel (light blue structure). Right: Number of inverse blebs observed throughout development on the distal and proximal halves of 17 embryos. p value results from paired Student’s t test.

**Supplementary Figure 3:**
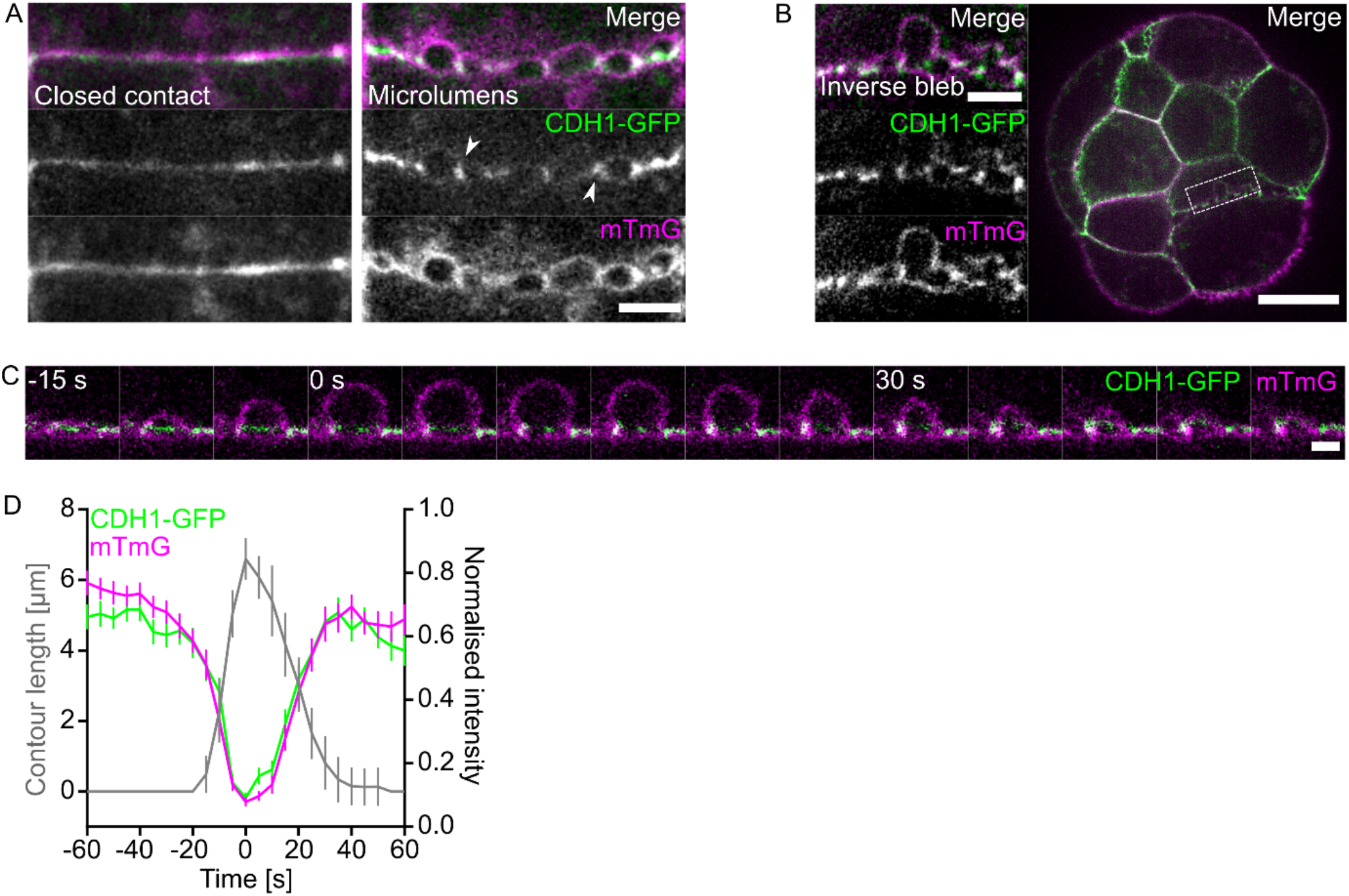
Dynamics of adhesion molecules in individual inverse blebs. A) Still images showing cell-cell contacts before and after the formation of microlumens with CDH1-GFP in green and mTmG in magenta. Arrowheads point at foci of CDH1 around microlumens. Scale bar, 5 μm. B) Equatorial plane of a 32-cell stage embryo with CDH1-GFP in green and mTmG in magenta (Movie 6). Scale bar, 20 μm. Dashed rectangle shows a cell-cell contact with an inverse bleb magnified on the left. Scale bar, 5 μm. C) Still images following the lifetime of an inverse bleb shown with mTmG in magenta and CDH1-GFP in green. Images taken every 5 s with time indicated relative to the maximal inverse bleb size. Scale bar, 2 μm. D) Mean contour length (grey), normalized CDH1-GFP (green) and mTmG (magenta) intensities of 31 inverse blebs from 7 embryos synchronized to their maximal extension. Error bars show SEM.

**Supplementary Figure 4:**
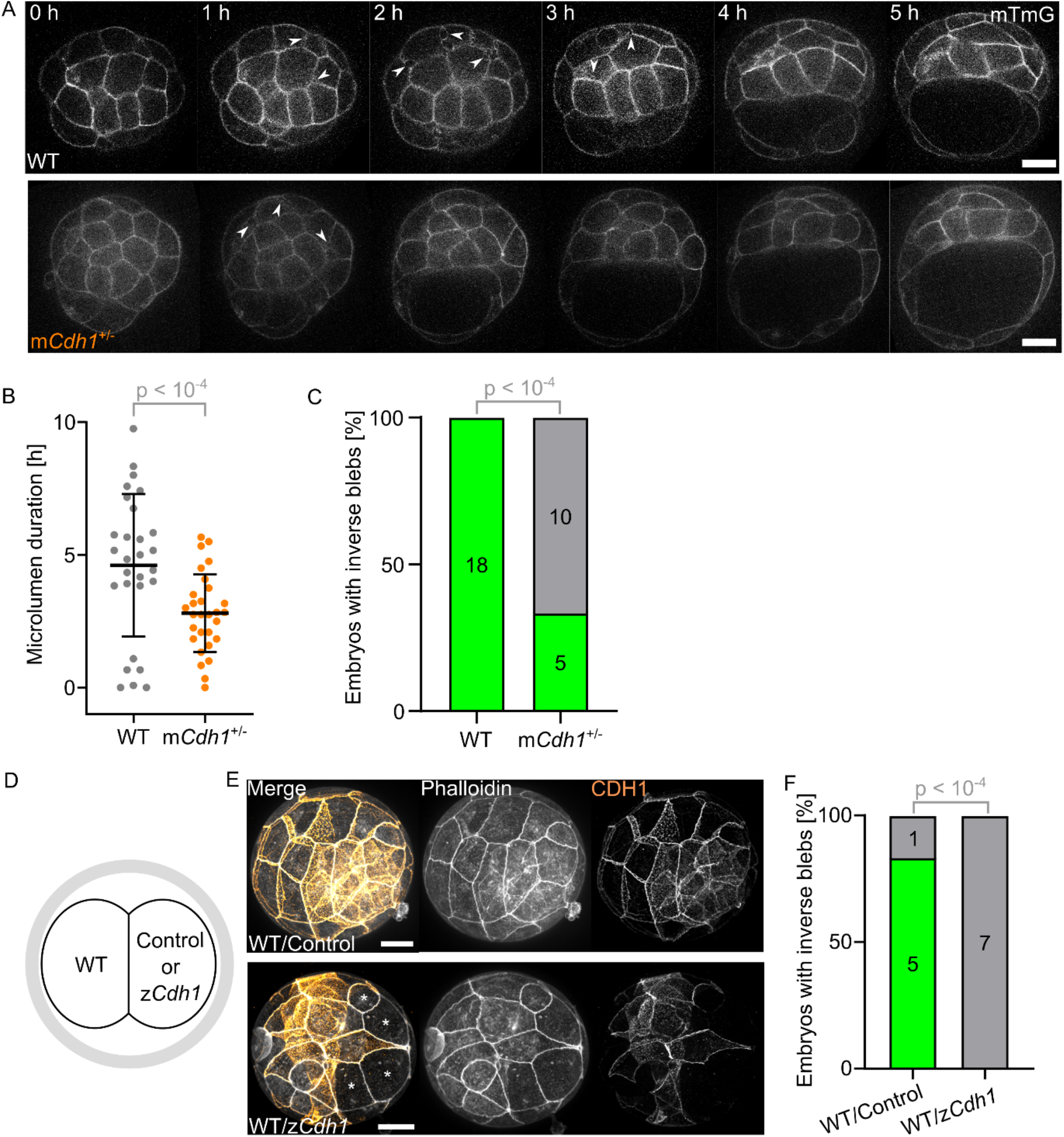
Global and local effects of adhesion loss on microlumens and inverse blebs. A) Still images of mTmG in WT (top) and m*Cdh1*^+/-^ (bottom) embryos during lumen formation (Movie 7). Arrows point at microlumens. Scale bars, 20 μm. B) Duration of microlumens in WT (n = 28) and m*Cdh1*^+/-^ (n = 29) embryos. Mean and SD are shown in black bracket. p value results from Welch’s t test. C) Proportion of WT (n = 18) and m*Cdh1*^+/-^ (n = 15) embryos showing inverse blebs. p value results from Fisher’s exact test. D) Schematic diagram of 2-cell stage embryo where one blastomere is microinjected with either CRISPR/Cas9 targeting *Cdh1* (z*Cdh1*) or Cas9 alone (Control). E) Immunostaining of embryos injected with Cas9 alone (Control) or Cas9 and CRISPR gRNA targeting *Cdh1* (z*Cdh1*) in one blastomere at the 2-cell stage with CDH1 shown in orange and actin in grey. Asterisks mark cells without CDH1. Scale bar, 20 μm. F) Proportion of embryos showing inverse blebs after injection with CRISPR/Cas9 targeting *Cdh1* (z*Cdh1*, n = 7 embryos) or Cas9 alone (Control, n = 6 embryos) in one blastomere at the 2-cell stage. p value results from Fisher’s exact test.

**Supplementary Figure 5:**
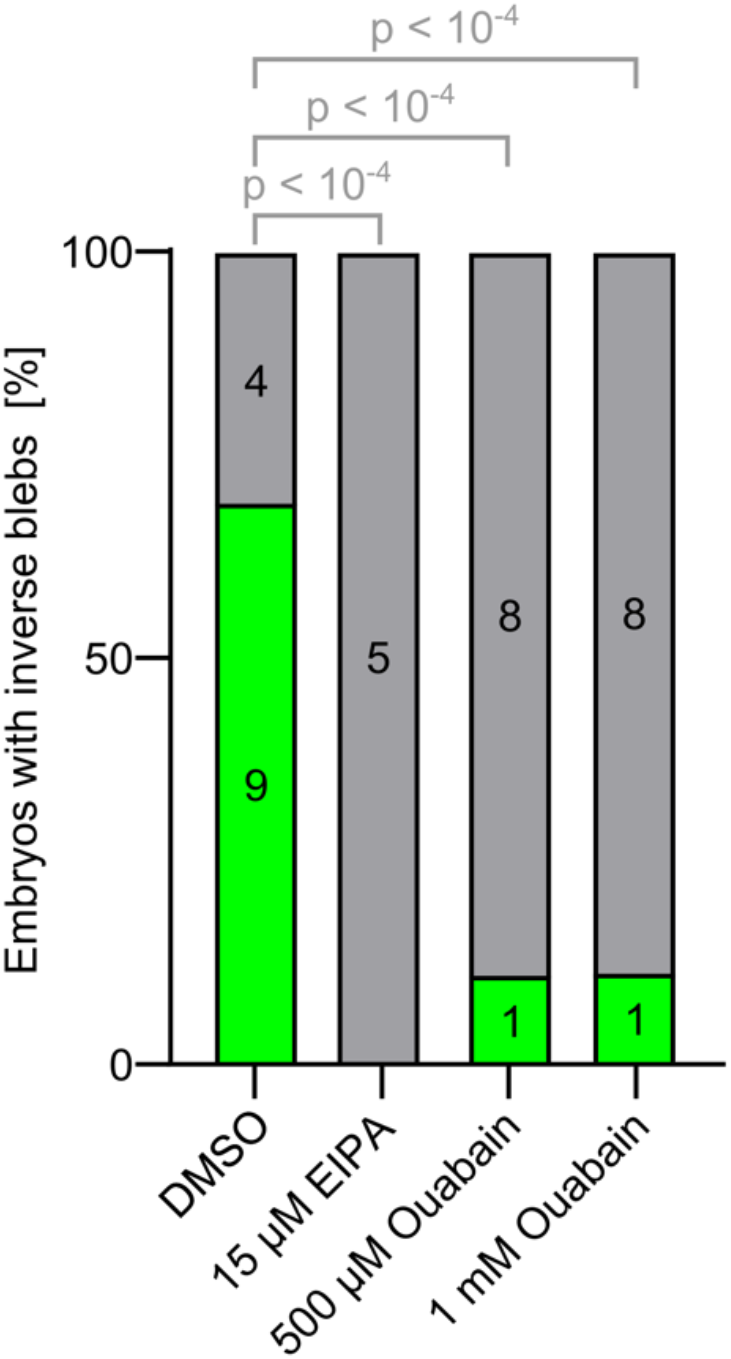
Inverse blebs require accumulation of luminal fluid. Proportion of embryos showing inverse blebs in DMSO (1:100 or 1:200, 13 embryos), EIPA (15 μM, n = 5 embryos) or, 500 μM (n = 9 embryos) or 1 mM (n = 9 embryos) Ouabain treatments during the 32-cell stage. p values result from Fisher’s exact test.

**Supplementary Figure 6:**
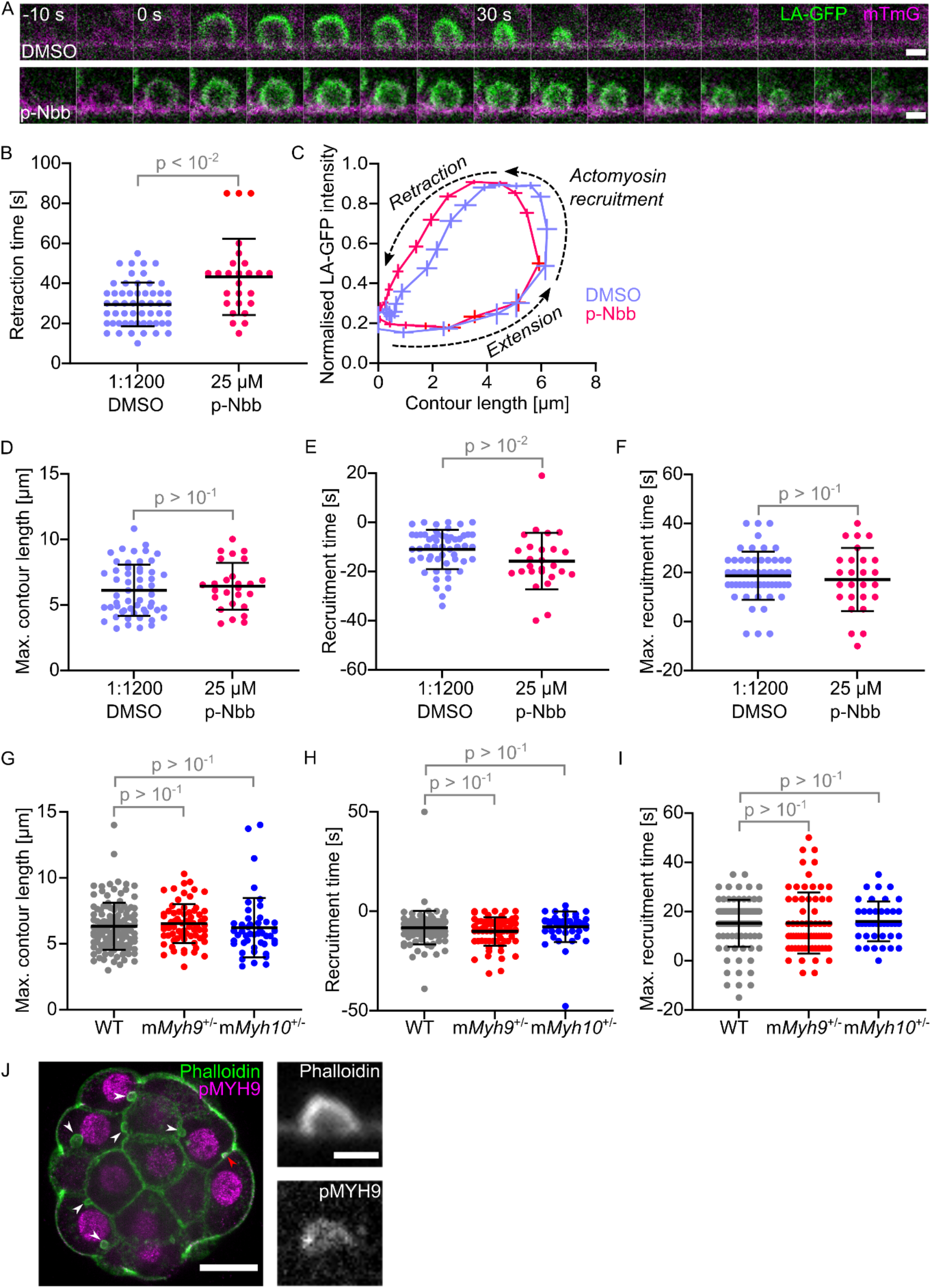
Dynamics of individual inverse blebs with impaired actomyosin contractility. A) Still images following the lifetime of an inverse blebs shown with LifeAct-GFP (LA-GFP) in green and mTmG in magenta in 1:1200 DMSO (top) or 25 μM para-Nitroblebbistatin (pNbb, bottom) media (Movie 10). Images taken every 5 s with time indicated relative to the maximal inverse bleb size. Scale bars, 2 μm. B) Duration of retraction of inverse blebs for embryos treated with 1:1200 DMSO (purple, n = 55 from 6 embryos) or 25 μM para-Nitroblebbistatin (violet, n = 26 from 6 embryos) media. Mean and SD are shown in black bracket. p value results from Welch’s t test. C) Mean normalized LA-GFP intensity as a function of mean contour length of inverse blebs synchronized to their maximal extension for embryos treated with 1:1200 DMSO (purple, n = 55 from 6 embryos) or 25 μM para-Nitroblebbistatin (violet, n = 26 from 6 embryos) media. Error bars show SEM. Arrows indicate the extension and retraction dynamics of inverse blebs. D-F) Maximal contour length, initial and maximal recruitment time of LA-GFP to inverse blebs of embryos treated with 1:1200 DMSO alone (purple, n = 55 blebs from 6 embryos) or with 25 μm para-Nitroblebbistatin (violet, n = 26 blebs from 6 embryos). Mean and SD are shown in black bracket. p values result from Welch’s t test. G-I) Maximal contour length, initial and maximal recruitment time of LA-GFP to inverse blebs of WT (grey, n = 100 blebs from 13 embryos), m*Myh9*^+/-^ (red, n = 67 blebs from 10 embryos) or m*Myh10*^+/-^ (blue, n = 41 blebs from 6 embryos) embryos. Mean and SD are shown in black bracket. p values result from ANOVA and Dunnett’s multiple comparisons tests. J) Left: immunostaining of 32-cell stage embryos showing Phalloidin (green) and the phosphorylated form of non-muscle myosin heavy chain IIA (pMYH9, magenta). Arrowheads point at fixed inverse blebs. Scale bar, 20 μm. Right: greyscale images on the right show separate signals of the inverse bleb pointed by the red arrowhead. Scale bar, 2 μm.

**Supplementary Figure 7:**
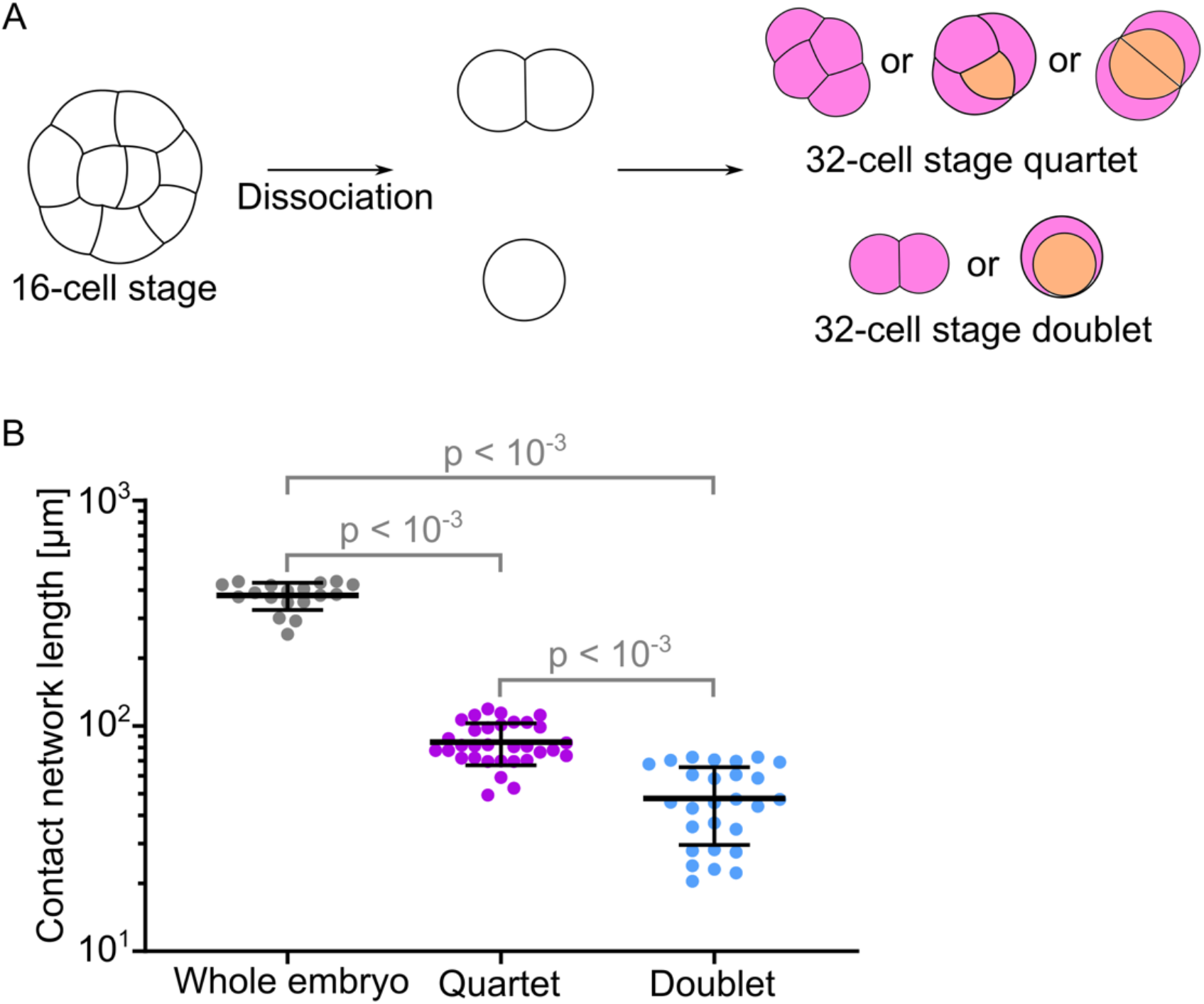
Geometry and topology of reduced embryos. A) Schematic diagrams of the dissociation process used to produce quartets and doublets of 32-cell stage blastomeres. A 16-cell stage embryo is dissociated into singlets and doublets which then divide into doublets and quartets respectively. Quartets can adopt 3 different configurations: from left to right, 4 outer cells, 3 outer and 1 inner cell or 2 outer and 2 inner cells, which have at least 2 multicellular microlumens. Doublets can consist of 2 outer cells or 1 outer and 1 inner cells, which do not have multicellular microlumen. B) Length of the network of cell-cell contact measured at the equatorial plane of whole embryos (n = 18), quartets (n = 32) or doublets (n = 27) shown with a logarithmic scale. p values result from ANOVA and Dunnett’s multiple comparisons tests.

## Movie legends

**Movie 1: Dynamic recruitment of actin during inverse blebs retraction**. Time lapse imaging at the equatorial plane of a 32-cell stage LifeAct-GFP (green) mTmG (magenta) embryo imaged every 5 s using spinning disk microscopy. Scale bar, 20 μm.

**Movie 2: Dynamic recruitment of non-muscle myosin II during inverse blebs retraction**. Time lapse imaging at the equatorial plane of a 32-cell stage MYH9-GFP (green) mTmG (magenta) embryo imaged every 5 s using spinning disk microscopy. Scale bar, 20 μm.

**Movie 3: Inverse blebs fill with intercellular fluid before flushing it back into the microlumen network**. Time lapse imaging at the equatorial plane of a 32-cell stage mTmG (grey) embryo incubated with Dextran-Alexa488 (cyan) at the 16-cell stage before sealing of tight junctions imaged every 5 s using spinning disk microscopy. Scale bar, 20 μm.

**Movie 4: Inverse bleb activity peaks before lumen expansion**. Nested time lapse imaging of a LifeAct-GFP embryo imaged every 10 s for 5 min repeated every 30 min using light sheet microscopy. Equatorial plane is shown on the left and maximum projection on the right. Developmental time is indicated relative to peak blebbing activity measured throughout the entire embryo volume. Scale bar, 20 μm.

**Movie 5: Inverse blebs coexist with microlumens throughout lumen formation**. Time lapse imaging at the equatorial plane of a LifeAct-GFP (green) mTmG (magenta) embryo imaged every minute using spinning disk microscopy. Scale bar, 20 μm.

**Movie 6: Dynamics of adhesion molecules during inverse blebs**. Time lapse imaging at the equatorial plane of a 32-cell stage CDH1-GFP (green) mTmG (magenta) embryo imaged every 5 s using spinning disk microscopy. Scale bar, 20 μm.

**Movie 7: m*Cdh1***^**+/-**^ **embryos form their lumen quicker than WT embryos**. Time lapse imaging of 32-cell stage mTmG WT and m*Cdh1*^+/-^ embryos imaged every 5 min using spinning disk microscopy. Embryos are imaged throughout their entire volume to synchronize them to the time of appearance of microlumens. The equatorial plane is shown. Scale bar, 20 μm.

**Movie 8: m*Cdh1***^**+/-**^ **embryos rarely form inverse blebs unlike WT embryos**. Time lapse imaging at the equatorial plane of 32-cell stage LifeAct-GFP (green) mTmG (magenta) WT and m*Cdh1*^+/-^ embryos imaged every minute using spinning disk microscopy. Embryos are synchronized to the time of appearance of microlumens at the equatorial plane. Scale bar, 20 μm.

**Movie 9: EIPA treatment acutely stops inverse blebs**. Time lapse imaging at the equatorial plane of a 32-cell stage LifeAct-GFP (green) mTmG (magenta) embryo imaged every 5 s using spinning disk microscopy. The embryo is first treated with 1:2500 DMSO containing medium (left), then with 20 μM EIPA for 15 min before the start of the time lapse (right). Scale bar, 20 μm.

**Movie 10: para-Nitroblebbistatin treatment acutely slows inverse bleb retraction**. Time lapse imaging at the equatorial plane of a LifeAct-GFP (green) mTmG (magenta) embryo imaged every 5 s using spinning disk microscopy. Treatment with 25 μM para-Nitroblebbistatin (p-Nbb) for 15 min before the start of the time lapse causes inverse blebs retraction to slow down compared to DMSO treatment. Scale bar, 20 μm.

**Movie 11: m*Myh9***^**+/-**^ **embryos retract inverse blebs slower than WT and m*Myh10***^**+/-**^ **embryos**. Time lapse imaging at the equatorial plane of WT, m*Myh9*^+/-^ or m*Myh10*^+/-^ LifeAct-GFP (green) mTmG (magenta) embryo imaged every 5 s using spinning disk microscopy. Scale bar, 20 μm.

**Movie 12: quartets of 32-cell stage blastomeres form a lumen rapidly**. Time lapse imaging at the equatorial plane of LifeAct-GFP (green) mTmG (magenta) quartet of 32-cell stage blastomeres imaged every min using spinning disk microscopy. Time is indicated relative to the appearance of microlumens at the equatorial plane. Scale bar, 20 μm.

**Movie 13: doublets of 32-cell stage blastomeres do not form a lumen rapidly**. Time lapse imaging at the equatorial plane of LifeAct-GFP (green) mTmG (magenta) doublet of 32-cell stage blastomeres imaged every min using spinning disk microscopy. Time is indicated relative to the appearance of microlumens at the equatorial plane. Scale bar, 20 μm.

## Methods

### Embryo work

All animal work is performed in the animal facility at the Institut Curie, with permission by the institutional veterinarian overseeing the operation (APAFIS #11054-2017082914226001 and APAFIS #39490-2022111819233999 v2). The animal facilities are operated according to international animal welfare rules.

Embryos are isolated from superovulated female mice mated with male mice. Superovulation of female mice is induced by intraperitoneal injection of 5 international units (IU) pregnant mare’s serum gonadotropin (PMSG, Ceva, Syncro-part), followed by intraperitoneal injection of 5 IU human chorionic gonadotropin (hCG, MSD Animal Health, Chorulon) 44-48 hours later. Embryos are recovered at E1.5 or E2.5 by flushing oviducts or/and uteri from plugged females with 37°C FHM (LifeGlobal, ZEHP-050 or Millipore, MR-122-D) using a modified syringe (Acufirm, 1400 LL 23).

Embryos are handled using an aspirator tube (Sigma, A5177-5EA) equipped with a glass pipette pulled from glass micropipettes (Blaubrand intraMark or Warner Instruments).

Embryos are placed in KSOM (LifeGlobal, ZEKS-050 or Millipore, MR-107-D) supplemented with 0.1 % BSA (Sigma, A3311) in 10 μL droplets covered in mineral oil (Sigma, M8410 or Acros Organics). Embryos are cultured in an incubator with a humidified atmosphere supplemented with 5% CO_2_ at 37°C.

To remove the zona zellucida (ZP), embryos are incubated for 45-60 s in pronase (Sigma, P8811). To dissociate blastomeres, embryos are incubated in EDTA containing Ca^2+^ free KSOM^42^ at the 16-cell stage for 3-5 minutes, followed by repeated aspiration into a smoothened glass capillary and dissociation into single 16-cell blastomeres or 16-cell blastomere doublets, which results, after development in KSOM, in the formation of 32-cell stage doublets or quartets, respectively.

For imaging, embryos are placed in 5 mm glass-bottom dishes (MatTek).

### Mouse lines

Mice are used from 5 weeks old on. To visualize plasma membranes and F-actin, *(Gt(ROSA)26Sor* ^*tm4(ACTB-tdTomato,-EGFP)Luo)*^*;Tg(CAG–EGFP)#Rows* mice^43,44^ are used. To visualize plasma membranes and MYH9, *(Gt(ROSA)26Sor* ^*tm4(ACTB-tdTomato,-EGFP)Luo*^*;Myh9* ^*tm8*.*1RSad*^ mice^43,45^ are used. To visualize plasma membranes and CDH1 *(Gt(ROSA)26Sor* ^*tm4(ACTB-tdTomato,-EGFP)Luo*^; *Cdh—GFP) mice*^*3,43*^ are used. To remove LoxP sites specifically in oocytes, *Zp3-cre (Tg(Zp3-cre)93Knw)* mice^46^ are used. To generate mMyh9^+/-^ embryos, *Myh9* ^*tm5RSad*^ mice^47^ are used to breed *Myh9* ^*tm5RSad/tm5RSad*^ ; *Zp3* ^*Cre/+*^ *females*^*46,47*^. To generate m*Myh10*^+/-^ embryos, *Myh10*^*tm7Rsad*^ mice are used to breed *Myh10*^*tm7Rsad/tm7Rsad*^; *Zp3* ^*Cre/+*^ *females*^*46,48*^. To generate m*Cdh1*^+/-^ embryos, *Cdh1*^*tm2kem*^ mice^49^ are used to breed *Cdh1*^*tm2kem/tm2kem*^ ; *Zp3*^*Cre/+*^ females^46,49^.

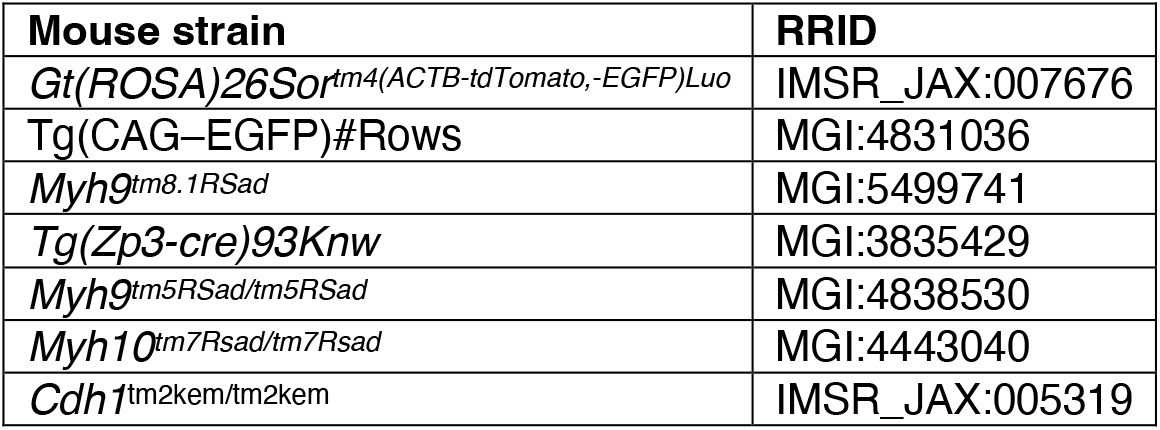

### Zygotic Cdh1 knockout in 2-cell embryos

To knockout *Cdh1*, we used the same approach as previously^50^. To target *Cdh1* a set of two gRNAs for each gene targeting close to the start codon are designed using the web tool CHOPCHOP^51^, namely GCGCGGAAAAGCTGCGGCAC and GCAGGAGCGCGGAAAAGCTG. For RNA transcription, the gRNA target sequences are cloned into pX458 (Addgene, 48138) as previously described^52^. Briefly, following digestion of the plasmid with BbsI, two annealed oligos encoding the gRNA target region, containing the 5’ overhang AAAC and the 3’ overhang CACCG are phosphorylated and cloned into pX458. The gRNA and its backbone are amplified by PCR with a fwd primer containing a T7 site. Following PCR purification RNA is transcribed using the MEGAshortscript T7 transcription kit (Invitrogen, AM1354). The RNA is then purified using the MEGAclear kit (Invitrogen, AM1908). Glass capillaries (Harvard Apparatus glass capillaries with 780 *μ*m inner diameter) are pulled using a needle puller and microforge to build a holding pipette and an injection needle. The resulting injection needles are filled with RNA/protein solution diluted in injection buffer (5mM Tris-HCl pH 7.4, 0.1 mM EDTA) to the following concentrations: To knock out *Cdh1*, 2-cell embryos are injected with 300 ng/*μ*L Cas9 protein (IDT, 1081058) and 80 ng/*μ*L of each gRNA diluted in injection buffer. To visualize injected cells, mRNA encoding lamin associated protein 2 fused to GFP (Lap2b-GFP) is added to the injection mix at 200 ng/μL^50^. Lap2b-GFP mRNA is transcribed using the mMESSAGE mMACHINE SP6 Kit (Invitrogen, AM1340) according to manufacturer’s instructions and resuspended in Rnase-free water.

The filled needle is positioned on a micromanipulator (Narishige MMO-4) and connected to a positive pressure pump (Eppendorf FemtoJet 4i). Control embryos were injected with Cas9 protein and Lap2b-GFP mRNA only.

Embryos are placed in FHM drops covered with mineral oil under Leica TL Led microscope. 2-cell embryos are injected while holding with holding pipette connected to a Micropump CellTram Oil. To verify the effectiveness of zygotic *Cdh1* knockout, immunostaining of chimeric embryo against CDH1 allows discerning the WT and knocked out halves of the embryos.

### Confocal microscopy

Live imaging of embryos is performed using a Viventis Microscopy LS1 Live microscope. Fluorescence excitation is achieved with a dual illumination scanned Gaussian beam light sheet of ∼1.1 μm full width 30% using a 488 nm laser. Signal is collected with a Nikon CFI75 Apo LWD 25x/1.1 objective and through a 488 nm long pass filter onto an Andor Zyla 4.2 Plus sCMOS camera. The microscope is equipped with an incubation chamber to keep the sample at 37°C and supply the atmosphere with 5% CO_2_. Embryos are sampled using a custom Python script every 10 s for 5 min, repeated every 30 min and the full volume of the embryo is acquired with a 76 μm z-stack with 2 μm step size.

Live imaging is performed using an inverted Zeiss Observer Z1 microscope with a CSU-X1 spinning disk unit (Yokogawa). Excitation is achieved using 488 and 561 nm laser lines through a 63x/1.2 C Apo Korr water immersion objective. Emission is collected through 525/50 and 595/50 band pass filters onto an ORCA-Flash 4.0 camera (C11440, Hamamatsu).

Alternatively, a Celldiscoverer 7 (Zeiss) with a CSU-X1 spinning disc unit (Yokogawa) is used. Excitation is achieved using 488 and 561 nm laser lines through a 50x/1.2 water immersion objective. Emission is collected through 514/30 and 592/25 band pass filters onto an ORCA-Flash 4.0 camera (C11440, Hamamatsu).

Both microscopes are equipped with an incubation chamber to keep the sample at 37°C and supply the atmosphere with 5% CO_2_.

In order to capture the dynamics of inverse blebs, WT embryos expressing LifeAct-GFP and mTmG or MYH9-GFP and mTmG, m*Myh9*^+/-^ or m*Myh10*^+/-^ embryos expressing LifeAct-GFP and mTmG are imaged every 5 s for 5 min at their equatorial plane, which is repeated every 1 h from late E2.5 to E3.5 using a custom Metamorph journal (Molecular Devices). A 60 μm z-stack with a 5 μm step size is acquired before every time lapse to discern TE and ICM cells.

To measure microlumen duration in WT and m*Cdh1*^+/-^ embryos, embryos expressing mTmG are imaged every 5 min for from E2.5 to E3.5, capturing a z-stack of 28 μm of the upper half of each embryo with a 4 μm step size.

In order to simultaneously track inverse bleb number and microlumens, WT or m*Cdh1*^+/-^ embryos, doublets or quartets are imaged at the equatorial plane every 1 min from before microlumen initiation until lumen formation is completed.

### Fluorescent dextran inclusion

3 kDa dextran coupled to Alexa Fluor 488 (Sigma, D34682) is added to KSOM at 0.1 g/L. Embryos are placed in labelled medium at the 16-cell stage before tight junctions fully seal.

### EIPA treatment

5-(N-Ethyl-N-isopropyl)-amiloride (EIPA) (Sigma, A3085) 50 mM DMSO stock is diluted to 20 μM in KSOM. A 1:2500 DMSO in KSOM dilution is used as control.

For acute EIPA treatment of embryos, embryos are cultured in the control medium and checked for the presence of inverse blebs. Embryos in the inverse bleb phase are imaged every 5 s for 10 min at their equatorial plane in the control medium. Embryos are then transferred to 20 μM EIPA medium and incubated for 15 min before being imaged in the same manner.

### para-Nitroblebbistatin treatment

Para-Nitroblebbistatin (Cayman Chemical) 50 mM DMSO stock is diluted to 25 μM in FHM without mineral oil. A 1:1200 DMSO in FHM dilution is used as control. Inverse blebbing embryos are either incubated in the control medium or para-Nitroblebbistatin medium for 15 min. Embryos are then imaged every 5 s for 10 to 30 min at their equatorial plane.

### Immunostaining

Embryos in the inverse bleb phase were fixed in 2% PFA (Euromedex, 2000-C) for 10 min at 37°C, washed in PBS, and permeabilized with 0.01% Triton X-100 (Euromedex, T8787) in PBS (PBT) at room temperature before being placed in blocking solution (PBT with 3% BSA) at 4°C for 2–4 h. Primary antibodies were applied in blocking solution at 4°C overnight. After washes in PBT at room temperature, embryos were incubated with secondary antibodies and phalloidin in blocking solution at room temperature for 1 h. Embryos were washed in PBT and imaged immediately after.

| Antibodies and dyes | Dilution | Provider | RRID |
| --- | --- | --- | --- |
| phospho-MYH9 (Ser1943) | 1:200 | Cell Signaling, 5026 | AB_10576567 |
| CDH1 | 1:500 | eBioscience, 14-3249-82 |  |
| Alexa Fluor 546 anti-mouse | 1:200 | Invitrogen, A11003 | AB_2534071 |
| Alexa Fluor Plus 488 anti-rabbit | 1:200 | Invitrogen, A32731 | AB_2633280 |
| Alexa Fluor 633 phalloidin | 1:200 | Invitrogen, A22284 |  |

## Data analysis

Data are analysed using Microsoft Excel and R. Data are plotted using Graphpad Prism.

### Inverse bleb count

The inverse bleb number in confocal and light-sheet time lapses is manually counted. A bleb is defined as a transient hemispherical actin-positive intrusion appearing at cell-cell contacts. When embryos are imaged every min continuously throughout lumen formation, the number of blebs is provided as such.

When embryos are sampled with nested time-lapses (brief acquisition every 5 or 10 s for 5 or 10 min repeated every 30-60min), the number of blebs is provided per min.

Inverse blebs are allocated to TE or ICM cells based on the position of the cells: TE cells contact the surface of the embryo while ICM cells are surrounded by other cells.

Inverse blebs are allocated into a distal and proximal half by determining the site of blastocoel formation in the equatorial plane and bisecting the embryo in two equal halves.

### Duration of the microlumen phase

Microlumen duration of embryos, quartets and doublets is determined as the time between the appearance of the first and disappearance of the last microlumens as detected visually.

### Dynamics of individual inverse blebs

Inverse bleb dynamics are measured using FIJI by tracing the outline of inverse blebs using a line width of 1 μm. Before and after inverse bleb formation and retraction, a line corresponding to the opening width of the inverse bleb is measured. For each inverse bleb, measurements are taken 60 s before and after reaching their maximum size (90 s after size maximum for para-Nitroblebbistatin treatment).

The time of maximal size of each inverse bleb is taken as a reference time point. Intensity values of LifeAct-GFP, MYH9-GFP, CDH1-GFP and mTmG are first normalized to 60 s before inverse bleb size maximum and then to their minimal and maximal values. The lifetime of an inverse bleb is taken as the time during which a visible membrane protrusion is seen extending at a cell-cell contact until the time it is no longer visible. The retraction time is taken as the time of inverse bleb duration from its maximal size to its disappearance.

Using R segmented package^53^, the LifeAct-GFP and MYH9-GFP recruitment times are calculated using iterative linear regression on the intensity data taken until the maximum value to find the breakpoint between the stable intensity and intensity increase.

### Blastocoel volume

Blastocoel volume is measured on full-volume light sheet z-stacks using the FIJI plugin LimeSeg^54^. The blastocoel is traced back from the expanded blastocyst and measured as soon as a lumen is present (smallest segmented lumen < 1 pL). Before that, blastocoel volume is assumed to be 0 pL.

### Statistics

Statistical test were performed using GraphPad Prism 9.5.1. Statistical significance is considered when p < 10^−2^.

The sample size was not predetermined and simply results from the repetition of experiments. No sample was excluded. No randomization method was used. The investigators were not blinded during experiments.

## Supplementary Note

In this note we briefly describe the minimal physical model for the formation and dynamics of intercellular fluid pockets situated at the interface of two adhering cells, henceforth referred to as fluid pockets. To capture the mechanics of these fluid pockets with volume *V*, surface area *A* and perimeter *P*, we propose an effective pseudopotential *W* where we consider the effects of surface tension *T*_0_ acting on the area, pressure build-up p and effect of adhesion at the periphery (Figure 1A). The surface tension seeks to shrink the fluid pockets while the pressure seeks to inflate them. Inside the fluid pockets, the cell membranes are detached but at the periphery there is enrichment of immobile adhesive bonds (within the timescale of ∼ 1 min), which can be thought of as elastic springs connecting the two plasma membranes (*1*). As more area detaches, more bonds detach and accumulate at the periphery (based on our experimental observations, also see (*2*)). The increased contraction from such elastic springs resist the increase in perimeter, while steric repulsion from increased number of adhesive bonds prefer to expand the perimeter. The balance of these two opposing effects set a preferred perimeter *P*_0_ and the mechanics can be captured by an effective perimeter elasticity with elastic coefficient k and rest perimeter *P*_0_ (see Appendix to the note). The pseudopotential W is then given by *W* = *T*_0_*A* + *k*(*P* − *P*_0_)^2^ − *pV*, and we also recast the rest perimeter as *P*_0_ = 2 π *R*_0_. For simplicity let us consider a spherical fluid pocket of radius R which yields

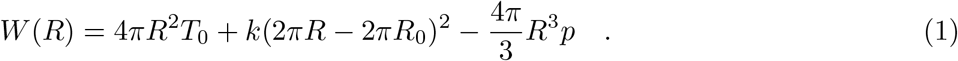

Note that here we consider an effective pressure *p* and do not explicitly take into account osmotic effects that may play a role at larger length and timescales (*3*). We nondimensionalize using the length scale *R*_0_ and tension *T*_0_ to define in relative units 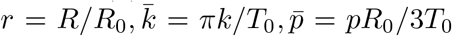. The nondimensional pseudopotential,

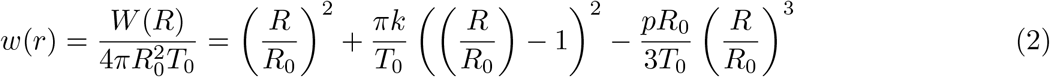

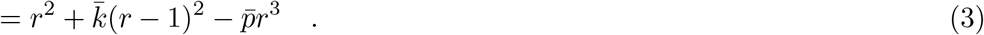

### Size of intercellular fluid pockets

The stable states of the system are captured by the minima of the pseudopotential *w*(*r*), which informs us of the mechanically stable size. The potential *w*(*r*) has a minima (*r*_*l*_) and a maxima (*r*_*h*_) at a finite *r* (see Fig.1A), given by the solution of d*w*/d*r* = 0 and is

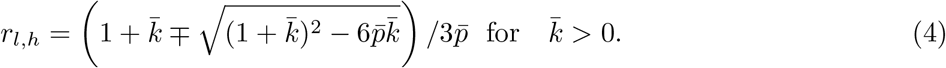

Near the minima *r*_*l*_ the fluid pockets relax to attain size *r*_*l*_, while beyond the maxima the fluid pockets grow indefinitely to maximal possible size (Figure 1B). In the *absence* of adhesion 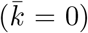, the minima *r*_*l*_ = 0. Hence, finite-sized fluid pockets are transient and should disappear at experimental timescales. Based on their initial size being larger or smaller than 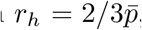, they will either grow very large (due to inflation) or shrink away (due to tension). This is similar to the nucleation mechanism of droplets and soap bubbles (*4*).

For 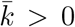, finite sized fluid pockets can exist at r_*l*_ given by Eq.(4) and for small 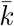 we find that the stable radius 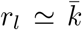. On the other hand, the maximum of the potential *w*(*r*) presents the barrier to nucleate a large-growing fluid pocket. This barrier is located at *r*_*h*_ and is given by Eq.(4). For small 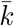 we can approximate 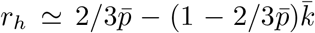. Due to fluctuations and continuous fluid pumping, a fluid pocket can cross over this potential barrier. Hence, for finite adhesion, stable fluid pockets of finite size and growing large fluid pockets of undetermined size can coexist. Note that the coexistence of stable fluid pockets and growing fluid pockets depends on the existence of distinct non-zero minimum *r*_*l*_ and maximum *r*_*h*_ and hence a finite barrier Δ*w*. As evident from Eq.(4), this is satisfied as long as 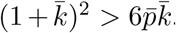. This allows us to derive a phase diagram as shown in Figure 2A. The propensity (or rate) to form such growing fluid pockets depends on the size of the potential barrier Δ*w* = *w*(*r*_*h*_) −*w*(*r*_*l*_). The lower the potential barrier, the higher is the probability to cross over the barrier. Δ*w* depends on 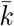 and 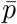 as

**Figure 1:**
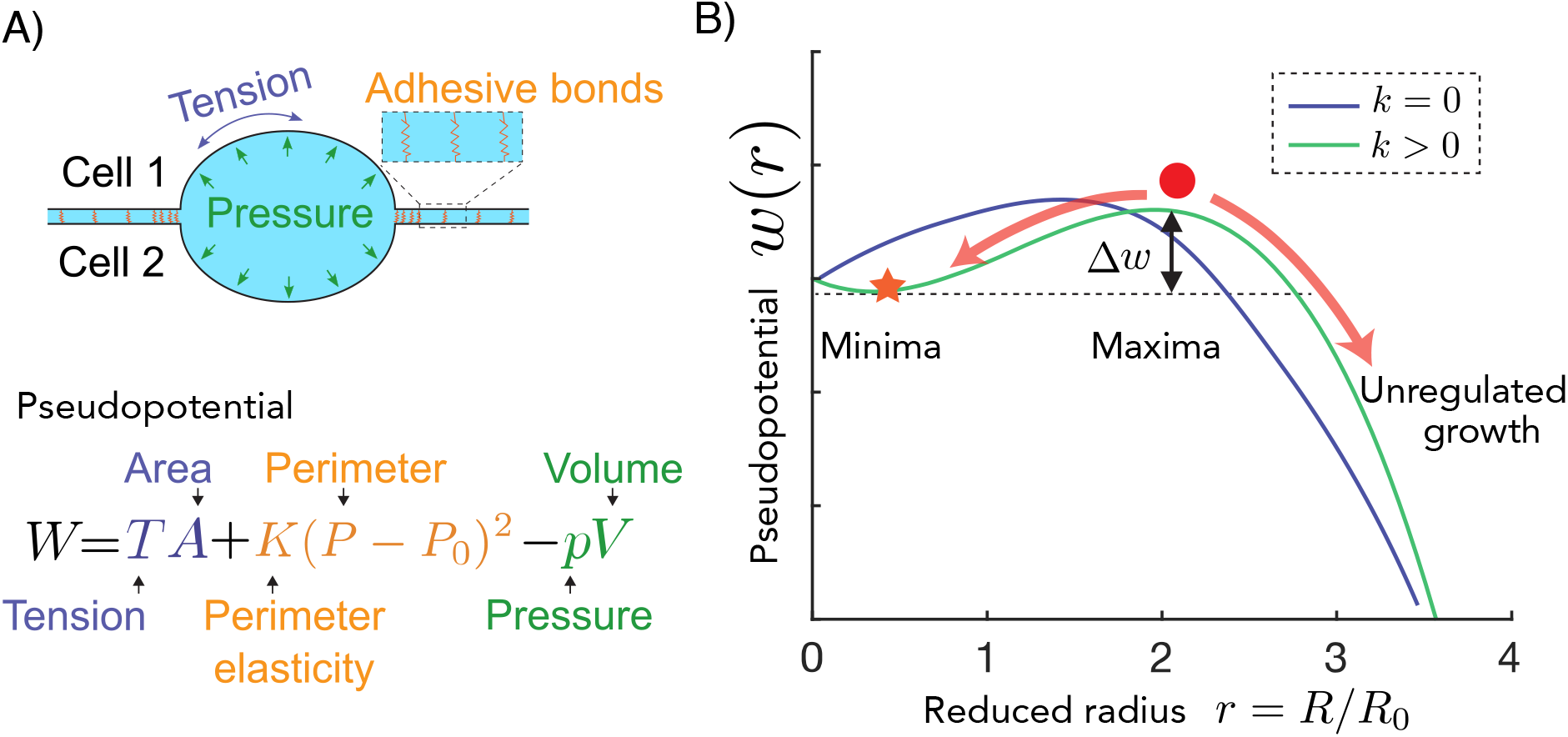
Mechanics of fluid pockets: A) Top: An intercellular fluid pocket situated between cell 1 and cell 2 is schematically depicted. Elastic adhesive bonds at the periphery of the fluid pocket are shown in the inset. Bottom: Energetic contributions from different mechanical parameters and geometric variables in the pseudopotential are depicted. B) Example normalised pseudopotential *w*(*r*) −*w*(0) are shown as a function of reduced radius *r* (see text). For *k* = 0 (violet), the minima lies at *r* = 0 implying no stable finite sized fluid pocket, while for *k* > 0 (green) a stable finite minima can be found (large orange star), implying coexistence of finite-sized fluid pockets with growing fluid pockets. The maxima is shown with a red dot. A potential barrier of height Δ*w* (the difference in *w* between the minima and maxima) is indicated for both cases.

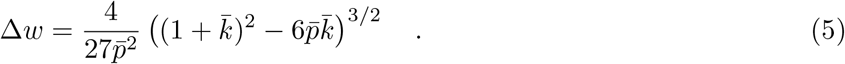

Note that Δ*w* is a decreasing function of relative pressure 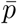 (see main text, Fig. 2G), which implies that build-up of pressure due to pumping lowers the barrier and helps the formation of growing fluid pockets. On the other hand, Δ*w* is an increasing function of relative perimeter elasticity 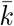, implying that adhesion inhibits barrier crossing (see maintext and Fig. 2F). Again, the barrier disappears when 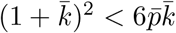, where the minima and the maxima fuse into an inflection point.

### Shape and symmetry of intercellular fluid pockets

We observe experimentally a characteristic difference in shape between the finite-sized fluid pockets and growing large fluid pockets. While the stable fluid pockets are typically symmetric and have a lens-like shape, the growing fluid pockets are asymmetric and resemble a spherical cap. To discuss whether a spontaneous symmetry breaking can take place in the course of the growth of the fluid pockets, we consider a spherical fluid pocket made of two hemispherical parts of radius *R*_1_ and *R*_2_ with identical tension *T*_0_ and perimeter elasticity *k*. The volumes 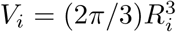, areas 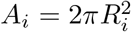 and perimeters *P*_*i*_ = 2π*R*_*i*_ for *i* = 1, 2. The mechanics can be captured with a pseudopotential similar to the previous section,

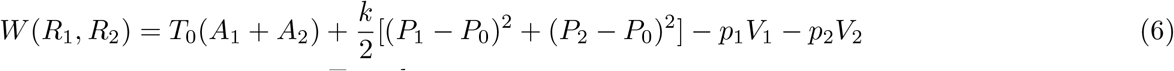

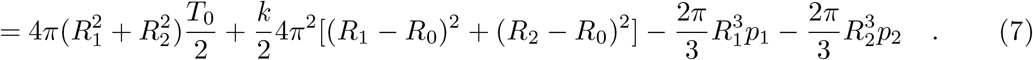

Note that here the hydrostatic pressure plays the role of a Lagrange multiplier (constraint force) to maintain separate but connected volumes. We can estimate the pressures *p*_1_ and *p*_2_ quasi-statically by seeking *dW*/*dR*_*j*_ = 0 for *j* = 1, 2 and find

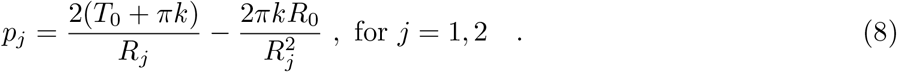

Note that for *k* = 0 we recover Laplace’s law. The volume balance in each of the hemispheres can be written as

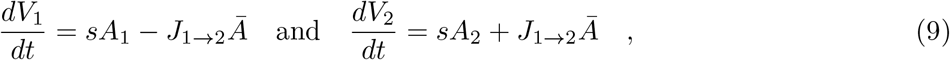

where *Ā* ≃ (*A*_1_ + *A*_2_)/2, s is pumping activity per unit area and *J*_1*→*2_ is a unit volume current from hemisphere 1 to hemisphere 2 due to change of shape. The current is described by a linear response law *J*_1*→*2_ = *μ*(*p*_1_ − *p*_2_) where the hydraulic mobility *μ* is a function of the sphere radii *R* and dynamic viscosity *η* given by *μ* ≃ *R*/3*η*. To study the symmetry breaking we study the dynamics of the relative volume difference *ν* = (*V*_1_ − *V*_2_)/(*V*_1_ + *V*_2_), given by

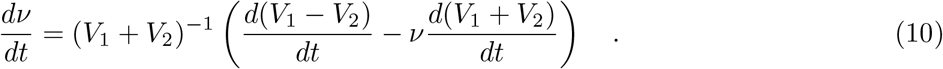

Note that *ν* = 0 is the symmetric configuration where both hemispheres are identical while *ν* = ±1 present the limiting cases where the entire volume is encapsulated either in hemisphere 1 or hemisphere 2. We consider a perturbation around the symmetric case (where *R*_1_ = *R*_2_ = *R*) in terms of the radii of the form *R*_1_ = *R*(1 + *ψ*) and *R*_2_ = *R*(1 − *ψ*) and perform a linear stability analysis. To linear order, the reduced volume difference *ν* = 3 *ψ*+ 𝒪 [*ψ*^3^] and the linearised dynamics can be captured by

**Figure 2:**
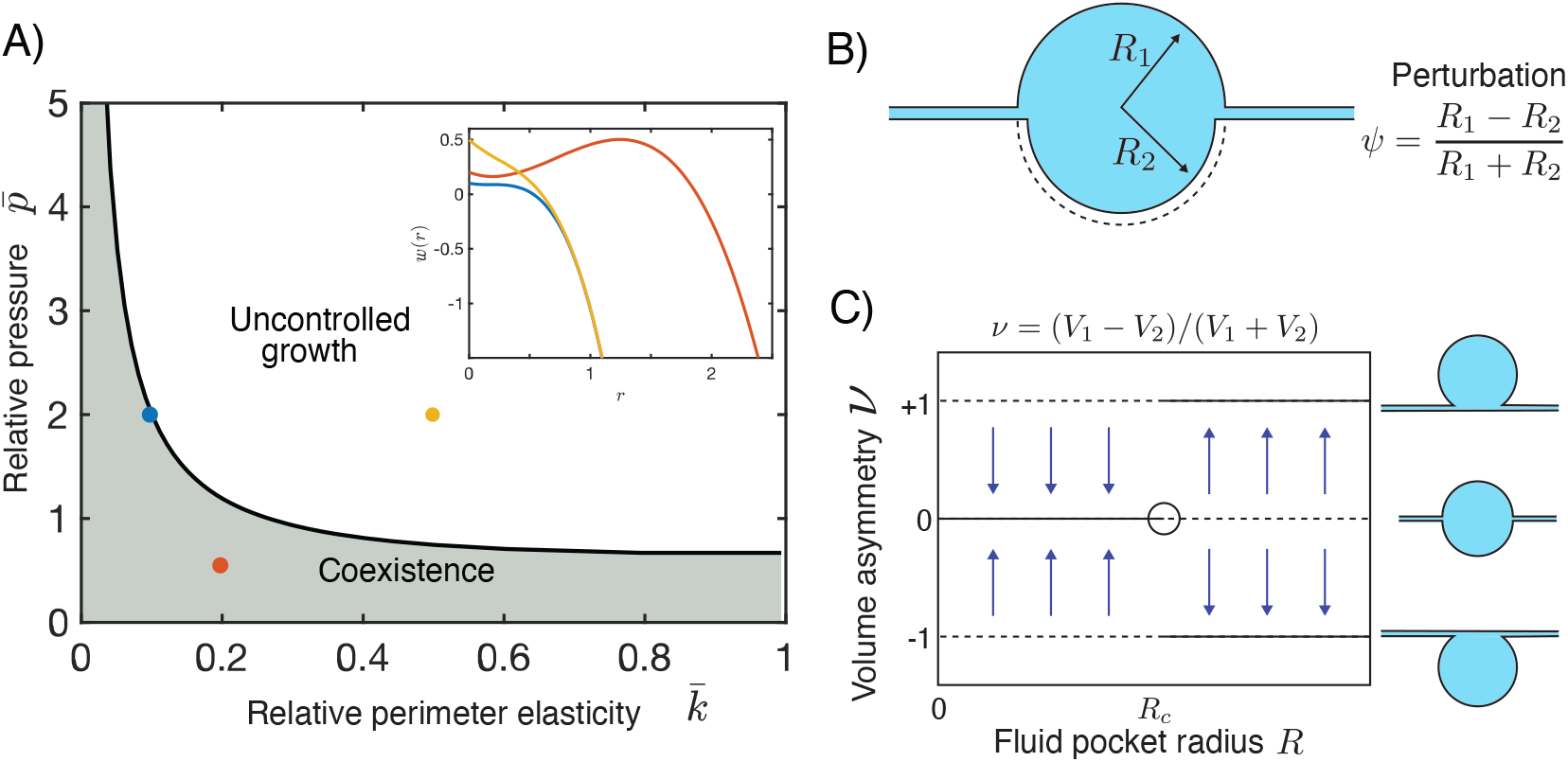
Phases and symmetries of fluid pockets: A) A phase diagram for coexistence of stable or growing fluid pockets is shown as function of relative adhesion 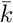 and relative pressure 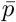. Shaded grey region indicates region of parameter space where coexistence is found. In the inset, pseudopotential *w*(*r*) is shown for colour coded parameter points. B) A shape perturbation scheme is depicted with two hemispheres of radius *R*_1_ > *R*_2_. C) A bifurcation diagram is shown for volume asymmetry *ν* as a function of system size/fluid pocket radius *R*. Below critical size *R*_*c*_, the symmetric shape *ν* = 0 is stable while above *R*_*c*_ the symmetric configuration *ν* = 0 is unstable and the system adopts the asymmetric state *ν* = ±1. The corresponding configurations are depicted on the right.

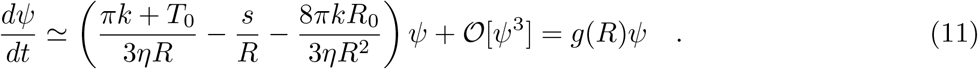

The asymmetry *ψ* grows when the growth rate *g*(*R*) is positive, leading to a symmetry breaking while for *g*(*R*) < 0 symmetry is restored. Beyond a critical size *R*_*c*_,

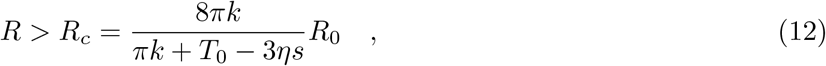

an instability (*5*) occurs where the symmetric state *ν* = *ψ* = 0 becomes unstable and relative differences between the two hemispheres diverge (see Fig. 2B). As a result we obtain an asymmetric configuration as depicted in Fig. 2B. On the other hand it implies that below *R*_*c*_, small asymmetries will decay and enforce symmetric shape of fluid pockets. Hence, fluid pockets during their growth can obtain their characteristic asymmetric shape via a growth induced mechanical instability leading to clear morphological distinction between symmetric microlumens and larger asymmetric inverse blebs.

### Retraction of inverse blebs

Inverse blebs grow fast and, following an enrichment of actomyosin, shrink away. Here we consider the effect of a time-modulated active tension *t*_*a*_ = *T*_*a*_/*T*_0_ in terms of actin enrichment Δ*ρ* (t) and study the resulting dynamics. The dynamics of the reduced radii *r* can be expressed as a relaxation process in potential *w*(*r*) with timescale *τ* and given as

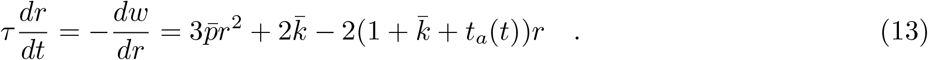

Surpassing the growth phase a transition from growing to shrinking can only occur for active tensions beyond a threshold given by

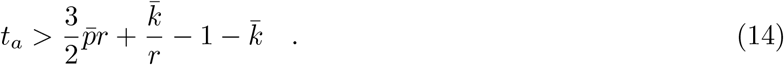

To satisfy this criteria, either an enrichment of actomyosin or a reduction in 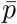 is necessary. Experimentally, we observe a sharp enrichment of actomyosin localisation in the retraction phase of the inverse bleb.

## Appendix

### Perimeter elasticity due to adhesive bonds

Consider a freshly nucleated spherical fluid pocket of radius *R*, where *N*_*a*_ number of adhesive bonds detached in the process of nucleation and accumulated at the periphery. If the surface density of adhesive sites prior to the detachment is *ρ*_*s*_, then N_*a*_ ≃2*π*R^2^*ρ*_*s*_. We can treat each of such adhesive bonds as an elastic spring with length *e*, rest length *e*_0_ and stiffness *k*_*e*_. Due to this enrichment, the density of adhesive bonds at the periphery *ρ*_*p*_ can be high. At high densities, interaction of molecules give rise to steric effects and cause repulsion. To capture the repulsive effects of such accumulation, we use the simplest energetic term Π/*ρ*_*p*_ that originates from a two dimensional steric pressure(similar to ideal gas). Note that the *ρ*_*p*_ = *N*_*a*_/2*πRd* = *Rρ*_*s*_/*d*, where *d* is the thickness of the peripheral region. The mechanics can be presented with an pseudopotential,

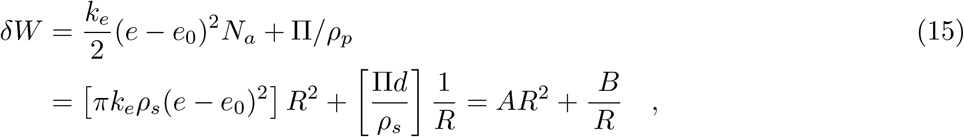

where 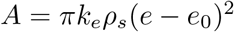 and *B* = Π*d*/*Ρ*_*s*_. The balance of these two effects sets a minima at

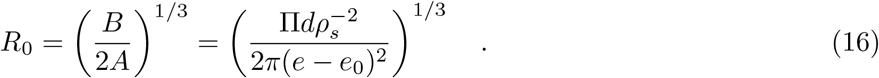

Around R_0_, we can perform an expansion up to the leading order term and we find an effective perimeter elasticity *k*(*P* − *P*_0_)^2^ with 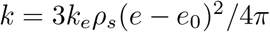,

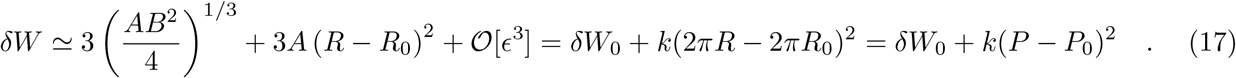

